# Creatine phosphagen system regulates energy metabolism and growth of trophoblasts

**DOI:** 10.64898/2026.05.04.722786

**Authors:** Nirvay Sah, Claire Zheng, Franny H. Stein, Walee B. Shaik, Maya B. Nassoura, Lauren M. Mole, Rachana Rajupalem, Donald Pizzo, Morgan Meads, Francesca Soncin

**Affiliations:** Department of Pathology, School of Medicine, University of California San Diego, 9500 Gilman drive, 92097, La Jolla, California, USA; Sanford Consortium for Regenerative Medicine, 2880 Torrey Pines Scenic drive, 92037, La Jolla, California, USA; Department of Poultry Science, University of Georgia, 120 D. W. Brooks Dr., 30602, Athens, GA, USA; Center for Precision Environmental Health, Baylor College of Medicine, 1 Baylor Plaza, 77030, Houston, Texas, USA

**Keywords:** Placenta, Trophoblast, Energy, Creatine, Creatine Kinase, Cyclocreatine

## Abstract

The creatine (Cr)-creatine kinase (CK)-phosphocreatine (PCr) system maintains ATP homeostasis in tissues with high energy demands, but its role in early human placental development remains poorly understood. We characterized Cr-CK-PCr system components across gestation and investigated its function in trophoblast development. Proteins involved in Cr biosynthesis, transport, and utilization were examined in first-trimester, second-trimester, and term human placentae (n=3-6/group). Components showed gestational age- and compartment-specific localization. The biosynthetic enzymes AGAT and GAMT localized predominantly to fetal villous mesenchyme and broadly across villous compartments, respectively. CKB was enriched in fetal mesenchyme, whereas CKMT1 was broadly expressed across trophoblast subtypes. The main Cr transporter SLC6A8 was abundant in syncytiotrophoblast early in gestation but shifted predominantly to fetal blood vessels at term. To determine functional importance, primary human trophoblast stem cells (hTSCs) were treated with a creatine analog, cyclocreatine (cCr; 0-40 mM), for 48 h. Treatment with 20 mM cCr reduced proliferation (P<0.001), decreased EGFR expression (P<0.05), induced cell-cycle arrest (P<0.0001), and increased apoptosis (P<0.0001). cCr also impaired mitochondrial and glycolytic function (P<0.01) with morphological evidence of mitochondrial stress. Consistently, treatment of primary human trophoblast organoids with cCr reduced organoid growth and EGFR expression (P<0.05). Together, these findings demonstrate that the Cr-CK-PCr system is dynamically expressed across human placental development and that disruption of Cr/PCr metabolism compromises trophoblast bioenergetics, proliferation, and survival, supporting its importance in placental development and providing a foundation for investigating its contribution to pregnancy disorders associated with altered placental energy metabolism.

**Summary sentence:** The creatine-creatine kinase-phosphocreatine system, whose components are dynamically expressed in a compartment-specific manner in human placenta across gestation, supports human trophoblast stem cell ATP metabolism, cell cycle and proliferation.

## Introduction

The placenta is a transient organ developed during pregnancy, which modulates the health of the mother and supports the growth of the fetus *in utero*. The fully formed human placenta is composed of villous trees made of epithelial trophoblast cells while the core is made of mesenchymal tissue containing fibroblasts, fetal blood vessels (endothelial cells and pericytes), and macrophages. Placental development precedes fetal development and as such, the placenta in early gestation mostly uses the nutrients from maternal endometrial secretions for its own growth and development [1]. While the embryo/fetus is developing *in utero*, the placenta performs a multitude of functions including cell biosynthesis, gas/nutrient transport, and hormone secretion to support fetal development. This makes the placenta a highly energy-demanding organ that must generate sufficient ATP to support its own bioenergetic and biosynthetic needs, as well as those of the growing fetus [2]. Trophoblast stem/progenitor cells, called cytotrophoblasts (CTB), which self-renew and differentiate into syncytiotrophoblasts (STB) or extravillous trophoblasts (EVT), drive early placental development through energy-intensive processes such as proliferation, migration, invasion, and fusion. Placental development continues well after the first trimester where the CTB continuously fuse to form STB, another process that requires ATP [1].

The primary pathways for ATP production include glycolysis and mitochondrial oxidative phosphorylation and the placenta is known to utilize both for energy production [3]. During high energy demands, homeostasis of ATP is tightly regulated to ensure cellular function and integrity. The creatine phosphagen system is the most studied system that buffers cellular ATP levels. Sources of creatine include diet and endogenous synthesis, mostly in the liver, kidneys and pancreas, mediated by the enzymes arginine:glycine amidinotransferase (AGAT) and guanidinoacetate methyl transferase (GAMT) [4]. Creatine is imported in target tissues/cells via the cognate transporter, SLC6A8, whereas SLC22A15 has been recently shown to control the cellular efflux of creatine [4,5]. In addition to these organs, studies suggest that on a cellular level, creatine can be synthesized locally by other organs that have high ATP requirements including brain [6], skeletal muscle [7], testes [8], gut [9], and placenta [10–12]. The ATP buffering capacity of the creatine phosphagen system is regulated by creatine kinase (CK), which catalyzes the reversible conversion of creatine (Cr) and ATP into phosphocreatine (PCr) and ADP. The two intracellular isoforms of CK, namely brain-type cytosolic CK (CKB) and ubiquitous mitochondrial CK (CKMT1), also acts together to shuttle ATP from the site of production (mitochondria) to the site of utilization (cytosol) [13], highlighting the “spatial energy buffering” role of the creatine phosphagen system.

The Cr-CK-PCr system has been implicated to maintain localized intracellular ATP recycling required for cell proliferation, migration, invasion, and maintenance of epithelial barrier as evidenced by studies across multiple species and systems [8,14–20]. Given that the cells of placenta alone use all of the aforementioned process for tissue development and functionality [21], the role of the Cr-CK-PCr system in placental bioenergetics and development could be critical. Previous studies in humans have confirmed the presence of Cr-CK-PCr system in term placenta [10] and studies from livestock models have showed the existence of this phosphagen system in first trimester placentae [22,23]. However, the exact role of the creatine phosphagen system in trophoblasts, the major epithelial cells of the placenta, and human placental development remain unknown. Multiple studies in humans have shown that disruption of the Cr-CK-PCr shuttle can be observed in placental dysfunctions including preeclampsia, fetal growth restriction, and diabetes mellitus [3,24–27] but the mechanistic role remains unclear. The placenta has greater growth and energetic dynamics in first trimester and any malfunction/formation during this early phase has a lasting influence on placental development and the health of mother and fetus [28]. Therefore, in this study, we first characterized the spatio-temporal expression of the components of the Cr-CK-PCr system across gestation in human placentae. Then, we determined the role of the creatine phosphagen system in trophoblast growth, using first trimester human trophoblast stem cells (hTSCs) and trophoblast organoids (TO) *in vitro*.

## Materials and methods

### Placental tissue samples

Bio-banked, formalin fixed paraffin embedded placenta tissues at the Center for Perinatal Discovery, University of California San Diego were used for immunostaining in this study. All bio-banked samples were collected under a UCSD Human Research Protections Program Committee Institutional Review Board-approved protocol; all patients gave informed consent following ethical and medical guidelines. Tissues samples were collected from early-first trimester (n=3, 6-8weeks GA), late-first trimester (n=3, 12-16 weeks), second trimester (n=3, 16-24 weeks), and term (n=3/male and female, 38-40 weeks GA) human pregnancies.

### Trophoblast stem cell culture

Primary human trophoblast stem cell lines 906p (male) and 1000p (female) derived from first trimester human placenta as described previously [29], between 18-25 passages, were used for this study. Primary hTSC were maintained in TSC complete medium containing Advanced DMEM/F-12 (Thermo Fisher, Cat# 12634010), supplemented with 1X B27/N2 (Thermo Fisher, Cat# 17502048, 17504044), 1X L-Glutamax (Thermo Fisher, Cat# 35050061), 0.05% BSA (Sigma Cat# A8412), 0.15 mM 1-thioglycerol (Sigma-Aldrich, Cat# M6145), 1% KSR (Thermo Fisher, Cat# 10-828-028), 2 μM CHIR99021 (Sigma Aldrich, Cat# SML1046-5MG), 0.5 μM A83–01 (Tocris, Cat# 2939), 1 mM SB43154 (Millipore, Cat# 616464), 0.8 mM VPA (Thermo Fisher, Cat# 676380), 5 μM Y27632 (SelleckChem, Cat# S1049), 100 ng/ml FGF2 (Biopioneer, Cat# HRP-0011), 50 ng/ml EGF (Millipore, Cat# 236-EG-01M), 50 ng/ml HGF (Stem Cell Technologies, Cat# 78019) and 20 ng/ml Noggin (R&D Systems, Cat# 6057-NG-025) [29,30]. hTSCs were grown in tissue culture-treated plates coated with 5 µg/mL collagen IV (Sigma, Cat #C0543-1VL) in incubator containing 5% CO_2_ and 21%O_2_ at a temperature of 37°C. They were passaged every three days using TrypLE, routinely checked for cell surface expression of EGFR and HLAG by flow cytometry, and tested for mycoplasma contamination. Cell counts were performed using the Countess automatic cell counter (Thermo Fisher).

### Trophoblast organoid derivation and culture, and cyclocreatine treatment

Trophoblast organoids (TOs) were derived from first trimester placental tissue (GA: 6 weeks, 3 days) and established in three-dimensional phenol red-free GFR Matrigel (Corning, Cat# 356231) domes for at least five passages prior to experiments. During passaging, TOs were broken down into single cell. Cell count and viability were then assessed, and cells were plated at a density of 20,000 cells per 15 µL Matrigel dome, with six-ten domes per condition. The domes containing TOs were then overlaid with 500 µL of trophoblast organoid media (t-TOM) [31]. The basal t-TOM medium comprised of Advanced DMEM/F12 (Thermo Fisher, Cat# 12634010), N2 (Thermo Fisher, Cat# 17502048), B27 (Thermo Fisher, Cat# 17504044), N-Acetyl-L-cysteine (Sigma Aldrich, Cat# A9165), Glutamax (Thermo Fisher, Cat# 35050061), nicotinamide (Sigma Aldrich, Cat# N0636), and FBS. Complete t-TOM medium included the addition of 500 nM A83-01 (Tocris, Cat# 2939), 1.5 µM CHIR99021 (Sigma Aldrich, Cat# SML1046-5MG), 50 ng/mL rhEGF (R&D Systems, Cat# 236-2EG), 50 ng/mL rhHGF (StemCell Technologies, Cat# 78019.2), 5 µM Y-27632 (SelleckChem, Cat# S1049), 80 ng/mL R-spondin-1 (PeproTech, Cat# 120-380-100UG), 100 ng/mL FGF-2 (Biopioneer, Cat# HRP-0011), and 2.5 µM PGE2 (SelleckChem, Cat# S3003).

### Cyclocreatine and creatine treatments

Fresh solutions of cyclocreatine (Sigma, Cat# 377627) and creatine (Sigma, Cat# C0780) were made for treatment of hTSCs and TOs and used within 3 days of preparation. A stock solution of 20 mM cCr was prepared in TSC or t-TOM medium at room temperature. Stock solutions were sterile filtered and further diluted in complete medium to reach desired concentrations. hTSC were plated at a density of 50k cells/well in a 12-well plate. Cells were allowed to attach for 12-16h before changing to fresh TSC medium containing either creatine or cyclocreatine. Treated cells were monitored for growth and collected for experimental assays after 48 hours of treatment. For TOs, treatment started five days after plating in fresh domes with 500 µL of t-TOM containing cyclocreatine for 6 days and media changed every other day. TOs were imaged daily under 4x magnification on an ECHO Rebel inverted microscope (ECHO) and collected after 6 days of treatment for analysis.

### Immunohistochemistry

Five-micron tissue sections were stained with antibodies to AGAT (1:2000; Sigma-Aldrich Cat# HPA026077), CKB (1:500; Abcam, Cat# ab92452), CKMT1a (1:4000; Biolegend, Cat# 867202), GAMT (1:200; Sigma-Aldrich, Cat# HPA051806), SLC6A8 (1:400; LSBio, Cat# LS-B13439), SLC6A13 (1:4000, Abcam, Cat# ab229815), and SLC22A15 (1:100; Sigma-Aldrich, Cat# HPA019785). Slides were stained on a Ventana Discovery Ultra (Ventana Medical Systems, Tucson, AZ, USA). Sections were deparaffinized by three cycles of heating to 72°C in the presence of Discovery Wash (Ventana, Cat# 07311079001) for 4 min. with a rinse between heating steps. Subsequently, antigen retrieval was performed using Discovery CC1 (Tris-EDTA; pH 8.6; Ventana, Cat# 06414575001) for either 40 min. total (5 cycles of 8 min. each with fresh CC1 for each incubation) at 100°C (AGAT, CKB, CKMT1, GAMT, SLC6A8, and SLC6A13) or 64 min (SLC22A15; 8 cycles of 8 min each). Endogenous peroxidase was quenched by incubation with inhibitor CM (Ventana, Cat# 760-159) for 8 minutes. The primary antibodies were incubated on the sections for 32 minutes at 37°C with the system mixing the samples every 4 minutes to ensure even distribution of the antibody. Primary antibodies were detected using the OmniMap system; an anti-rabbit (Ventana, Cat# 05266548001) or anti-mouse (Ventana, Cat# 5266556001) secondary coupled to an HRP polymer. Antibody presence was visualized used the Ventana Discovery ChromoMap DAB kit (Ventana, Cat# 05266645001) as a chromogen followed by hematoxylin II (Ventana, Cat# 05277965001) for 4 min. followed by a 4 min. treatment with bluing reagent (Ventana, Cat# 05266769001) as a counterstain. Slides were rinsed, dehydrated through alcohol and xylene and coverslipped. Slides were imaged using Olympus light microscope.

### Immunofluorescence Staining

Five-micron tissue sections were stained sequentially with various antibodies or in combinations (GAMT, CKB, SLC6A8/CD31; Frataxin/CKMT1) using the tyramide signal amplification system (TSA; ThermoFisher). Slides were stained on a Ventana Discovery Ultra (Ventana Medical Systems, Tucson, AZ, USA). Antigen retrieval was performed using CC1 for 40 minutes at 100°C. The first antibody in the dual staining was applied for 32 min at 37°C and used the appropriate species secondary antibody (Ventana) followed by fluorescent labeling with TSA-Alexa 488 according to the manufacturer’s instructions adapted for use on the Ventana. The antibodies were fully denatured, inactivated, and removed from the tissue by treatment in CC2 (pH 6.5; Ventana) for 20 min at 100°C. Subsequently, the second antibody was applied and detected by the species-appropriate secondary (Ventana) followed by fluorescent labeling with TSA-Alexa 594. Slides with no primary antibodies (no primary control) were stained simultaneously and used as negative controls (Supplemental Fig 1). After staining, sections were rinsed, Prolong gold antifade with DAPI was applied (Vectorlabs) and coverslipped. Slides were imaged on Echo Revolve fluorescent microscope.

For immunofluorescence staining of hTSCs, cells were fixed in culture plate with 4% paraformaldehyde for 20 minutes. Cells were then washed with 1X PBS three times and permeabilized with 0.1% Triton X for 10 min. at room temperature. Cells were then incubated with blocking buffer (5% BSA/0.05% Triton X) for 1h at room temperature, washed, and incubated with primary antibody overnight. The next day, cells were washed with 1X PBS and incubated with a suitable fluorophore conjugated secondary antibody and DAPI for 2h at room temperature in dark. Cells were then washed again and coverslipped for imaging. Images were taken using Echo Revolve fluorescent microscope.

### Transmission electron microscopy

For epoxy resin embedding, cells were fixed with 2.5% glutaraldehyde in 0.1 M sodium cacodylate buffer pH 7.4 for 5 minutes at room temperature and for 2h on ice, then pelleted in 10% gelatin. After washing with 0.1 M cacodylate buffer, pellets were postfixed in 1% OsO4 in 0.1 M cacodylate buffer for 1h on ice. Cells were stained all at once with 2% uranyl acetate for 1h on ice, following which they were dehydrated in graded series of ethanol (50-100%) while on ice. Cells were then subjected to 1 wash with 100% ethanol and 2 washes with acetone (10 minutes each) and embedded with Durcupan. Sections were cut at 60 nm on a Leica UCT ultramicrotome and picked up on 300 mesh copper grids. Sections were post-stained with 2% uranyl acetate for 5 minutes and Sato’s lead stain for 1 minute. Images were acquired with a transmission electron microscope Tecnai Spirit G2 Biotwin, operated at 80KeV and equipped with a bottom-mounted 4kx4k camera FEI Eagle. Morphometry analysis of mitochondria was performed in ImageJ for 100 mitochondria from 3-4 randomly selected cells of three technical replicates from each treatment group.

### Cell cycle and Apoptosis assay

Cell cycle analysis was performed using Click-iT EdU cell proliferation assay kit for imaging, Alexa Fluor 488 dye (ThermoFisher Scientific, Cat# C10337) as per manufacturer’s recommendations. After 48h of culture in cyclocreatine media, hTSCs were incubated with 10 µM of EdU or without EdU (used as negative control) for 2h in cell culture incubator. Cells were then dissociated with TrypLE and fixed in 100 µL of Click-iT fixative for 15 minutes at room temperature protected from light. Cells were then washed in 1%BSA in PBS and incubated for 15 minutes following resuspension in 100 µL of 1x Click-iT saponin-based permeabilization and wash reagent. EdU incorporation was detected by incubating the cells for 30 minutes at room temperature in Click-iT reaction cocktail as detailed in manufacturer’s protocol. Cells were washed in PBS and further incubated briefly in DAPI before performing flow cytometry analysis.

The cell apoptosis assay was performed using Dead cell apoptosis kits with Annexin V FITC and Propidium Iodide for flow cytometry (ThermoFisher Scientific, Cat# V1324) following manufacturer’s recommendations. hTSCs treated with cyclocreatine for 48h were dissociated from the plates with TrypLE and washed in cold PBS. Cells were resuspended in 100 uL of 1X annexin-binding buffer at a maximum concentration of 1 million cells/mL. FITC Annexin V (5 µL) and propidium iodide (1 µL) were then added to the cell suspension and incubated for 15 minutes at room temperature. Then, cells were washed, resuspended in 300 µL of 1X annexin-binding buffer and immediately used for flow cytometry analysis. One replicate of each treatment remained unstained for Annexin V and PI and were used as negative controls for the assay.

### Flow cytometry analysis

Cyclocreatine-treated cells were dissociated from culture plate with TrypLE and washed in FACS buffer consisting of 2% BSA and 0.03% sodium azide in PBS. Cells were then incubated either with APC conjugated-EGFR (Biolegend, Cat# 352906) and HLAG (Exbio, Cat# 1P-292-C100) antibodies or their isotype controls (Biolegend, APC mouse IgG1 Cat# 400122 and PE mouse IgG1 Cat# 400112), for 20 minutes at 4°C protected from light. Cells were then washed twice in PBS and fixed in 2% paraformaldehyde for analysis on the flow cytometer. All flow cytometry analysis in this study were performed using the FACSCanto or LSRFortessa X-20 (BD Biosciences) flow cytometer analyzers and further processed using FlowJo.

### Seahorse Mito-stress test and glycolytic stress test

Energy metabolism in cyclocreatine-treated hTSCs was measured using the Seahorse assays in 96-well plate at a seeding density of 10,000 cells/well. Cells were seeding in TSC complete medium and allowed to attach to the collagen IV-coated 96-well plate overnight followed by replacing with cyclocreatine medium. After 48h of culture, the cells were starved of media and the mito- or glycolytic stress test was performed according to manufacturers’ guidelines (Agilent Seahorse). Briefly, mitochondrial respiration was assessed using mitochondrial stress test by measuring oxygen consumption rate (OCR). Following baseline measurements, oligomycin was injected to inhibit ATP synthase, allowing determination of ATP-linked respiration and proton leak. FCCP was subsequently added to uncouple mitochondrial respiration and drive maximal electron transport, enabling assessment of maximal respiratory capacity and spare respiratory capacity. Finally, rotenone and antimycin A were injected to inhibit complexes I and III, respectively, to quantify non-mitochondrial respiration. Additionally, glycolytic function was evaluated using the glycolytic stress test by measuring extracellular acidification rate (ECAR). After baseline measurements, glucose was injected to initiate glycolysis and determine basal glycolytic activity. Oligomycin was then added to inhibit mitochondrial ATP production, thereby shifting energy dependence to glycolysis and revealing glycolytic capacity. Lastly, 2-deoxy-D-glucose was injected to inhibit hexokinase, allowing confirmation of glycolysis-dependent acidification and calculation of non-glycolytic acidification. Five technical replicates per treatment conditions (0, 1, 10, and 20mM cyclocreatine) were used for the assays.

### RNA extraction, cDNA synthesis, and quantitative real-time PCR

Total RNA was extracted from cells using NucleoSpin RNA columns (Macherey-Nagel, Cat# 740955.250) following manufacturers protocol. Cells were lysed in 350 µL of Buffer RA1 and stored at -20C until used for RNA isolation. Frozen cell lysates were thawed on ice, vortexed and the lysates were passed through the NucleoSpin Filter. The flow through was mixed with equal volume of 70% ethanol and the total volume was loaded on to RNA column followed by centrifugation at 11,000g for 30 seconds to allow RNA binding. The column was then desalted with MDB buffer and incubated with DNase for 15 minutes at room temperature to remove genomic DNA. The column was washed with buffers RAW2, RA3 and dried. RNA was eluted in 30 µL of nuclease free water. Concentration of RNA was measured on a NanoDrop 2000C spectrophotometer (Thermo Fisher). cDNA was synthesized using 100-300 ng of total RNA using PrimeScript RT kit (TaKaRa Bio Cat# RR037A). cDNA samples were diluted to obtain a working stock of 3 ng/µL to be used for qPCR assay. Primers for qPCR were designed using Primer-BLAST (NIH NCBI) and are provided in Supplementary Table S1. Real time quantitative PCR was performed in a 3 µL reaction volume containing 1 µL of cDNA, and 2µL of master mix composed of TB Green Premix Ex Taq (TaKaRa Bio, Cat# RR420A), ROXII, and a 12.5 μM mix of forward and reverse primers. The PCR was programmed for 40 cycles of 95°C for 5 seconds and 60°C for 34 seconds on QuantStudio 5 thermal cycler (Applied Biosystems). Primer specificity was confirmed using melt curves. Differential gene expression was performed using ΔΔCT method, normalized to 18S rRNA.

### ATP assay

Intracellular concentration of ATP in the hTSCs was quantified using guidelines provided by ATP Detection Assay Kit-Luminescence (Cayman Chemical, Cat#700410). After 48h of cyclocreatine treatment, hTSCs were dissociated from the plate and cell count was noted. Cells were homogenized in 1X ATP detection sample buffer and stored at -20C until analyzed. ATP assay was performed in 100 μL reaction volume containing 100 μL of reaction mixture (ATP detection assay buffer, D-Luciferin, and Luciferase) and 10 μL of samples. Samples were read on Tecan infinite M1000 pro microplate reader. Concentrations of ATP in samples were quantified using the equation of the standard ATP linear curve. Assay was performed with 5 technical replicates.

### Creatine Kinase activity assay

The activity of CK was measured using a colorimetric assay kit (Abcam, Cat# ab155901) according to the manufacturer’s protocol. Briefly, cyclocreatine-treated cells were resuspended in ice cold Assay Buffer. Assay was performed using 50 μL of diluted sample and 50 μL of reaction mix consisting of CK enzyme mix, developer solution, ATP, and creatine. Optical density (OD) of samples was read on Tecan infinite M1000 pro microplate reader in a kinetic mode every 2 minutes for 40 minutes at 37°C. Concentrations of NADH in samples were quantified by using the difference in OD values in the linear range. Final activity of CK in the samples were calculated by dividing the sample NADH content by the reaction time and volume of sample used times the dilution factor. Assay was performed with 5 technical replicates.

### Statistical analysis

Statistical analyses were performed using GraphPad Prism v10. Mean comparisons of two groups were done using the Student’s t-test, whereas statistical comparison of three or more groups was performed using one-way ANOVA followed by post-hoc Tukey’s test or Šídák’s multiple comparisons test for repeated measures. P values <0.05 were considered significant.

## Results

### Proteins involved in biosynthesis and transport of creatine are expressed in a compartment-specific manner in the placenta

We performed immunohistochemistry for proteins associated with the biosynthesis (AGAT and GAMT) and transport (SLC6A8) of creatine in the human placenta across gestation (Fig 1, Supplemental Fig. 1). AGAT, involved in production of the creatine precursor, guanidinoacetate (GA), localized predominantly to the villous mesenchymal compartment, including the fetal villous stromal cells and vascular endothelial cells (Fig 1A). This cellular localization pattern was conserved in the placenta across gestation, with the trophoblast compartment lacking expression of AGAT. On the other hand, GAMT, responsible for methylation of guanidinoacetate to creatine, was mostly localized in the trophoblast compartment including the CTB, cell column (precursors of EVT), and STB across gestation (Fig 1B and Supplemental Fig 2A). GAMT also appeared to be expressed in villous stromal and endothelial cells, but with much lower immunoreactivity than that in the trophoblast villous compartment (Fig 1B and Supplemental Fig 2A). Based on the IHC staining, the localization of the creatine transporter, SLC6A8, appeared to be dynamic across gestation; i) in first trimester, its expression was detected mostly in the villous compartment, particularly at the apical surface of STB, ii) in the second trimester, SLC6A8 localized to both the villous trophoblasts as well as endothelial cells, and iii) at term, a strong immunoreactivity for SLC6A8 was observed in the blood vessels and moderate immunoreactivity in fetal villous stromal compartment (Fig 1C). Immunofluorescence staining confirmed the loss of SLC6A8 expression from villous trophoblast with progression of pregnancy (Fig 1D). In contrast to the anticipated endothelial localization, the co-staining of placentae with SLC6A8 and the endothelial marker, CD31, revealed SLC6A8 signal mostly expressed in few circulating immune cells within the fetal capillaries (Fig 1D). The GA importer, SLC6A13, was consistently expressed in both the placental trophoblast and mesenchyme compartment with localization in CTB, STB, stromal and endothelial cells across gestation (Supplemental Fig 3A). SLC22A15, the transporter controlling efflux of creatine, was localized predominantly to the fetal villous endothelial cells across gestation suggesting potential export of creatine between placental and fetal tissues (Supplemental Fig 3B). These immunolocalization results suggest human placenta can produce creatine locally throughout gestation with co-operation between the villous mesenchymal and trophoblast compartments.

**Figure 1:**
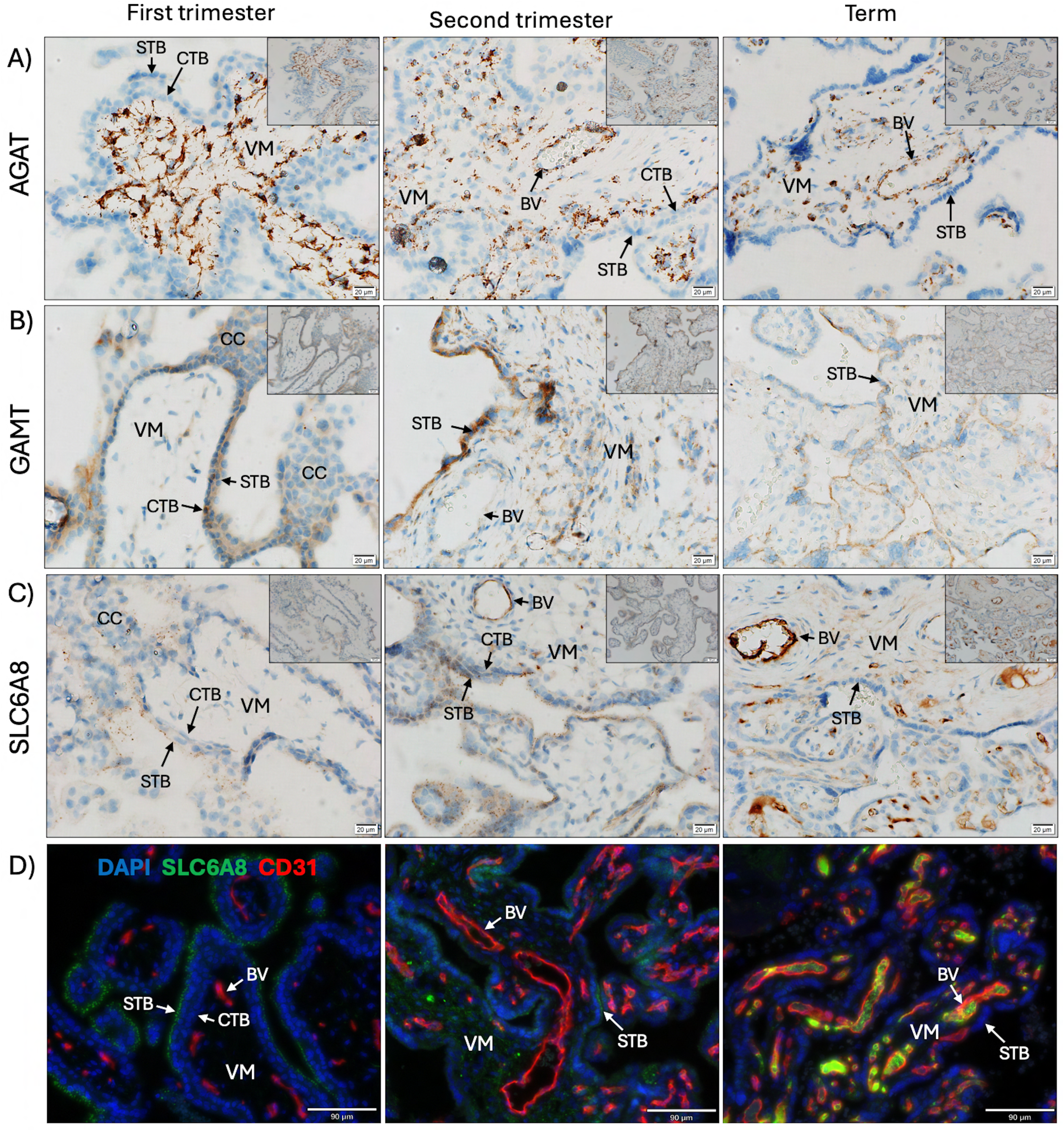
Immunolocalization of proteins associated with biosynthesis and transport of creatine in human placenta across gestation. A) AGAT localized to primarily to the placental villous mesenchymal (VM) compartment only throughout gestation. B) GAMT expression was observed mainly in the placental villous cytotrophoblast (CTB) and syncytiotrophoblast (STB) across gestation. C and D) Creatine transporter, SLC6A8, localized to the villous trophoblast in first trimester but shifted to blood vessels (BV) at term. Main images of IHC were taken at 40x (scale bar 20µm), and the insets were images at 20x (scale bar 50µm). Scale bar of the immunofluorescence staining are 90µm. Images are representative of first trimester (n=6), second trimester (n=3) and term (n=6) placentae.

### Creatine Kinases are dynamically expressed in the trophoblast compartment of the placenta

We performed immunohistochemistry for creatine kinase isoforms that differ in cellular localization with one predominantly cytoplasmic (CKB) and the other mitochondrial (CKMT1). The localization of the cytosolic creatine kinase protein (CKB) in placenta appeared to be dynamically altered across gestation. In first trimester, expression of CKB was observed in villous CTB and STB whereas, in the second trimester, its immunoreactivity was seen in both villous trophoblasts (CTB and STB) as well as villous mesenchyme including stromal and endothelial cells (Fig 2A). On the other hand, in the term placenta, CKB localized mostly to the endothelial cell and pericytes surrounding the blood vessels, with almost no immunoreactivity in CTB or STB (Fig 2A and Supplemental Fig 2B). In contrast, the mitochondrial creatine kinase, CKMT1, was consistently and exclusively localized to the placental trophoblast compartment including the CTB, STB, and cell column (Fig 2B) irrespective of gestational age.

**Figure 2:**
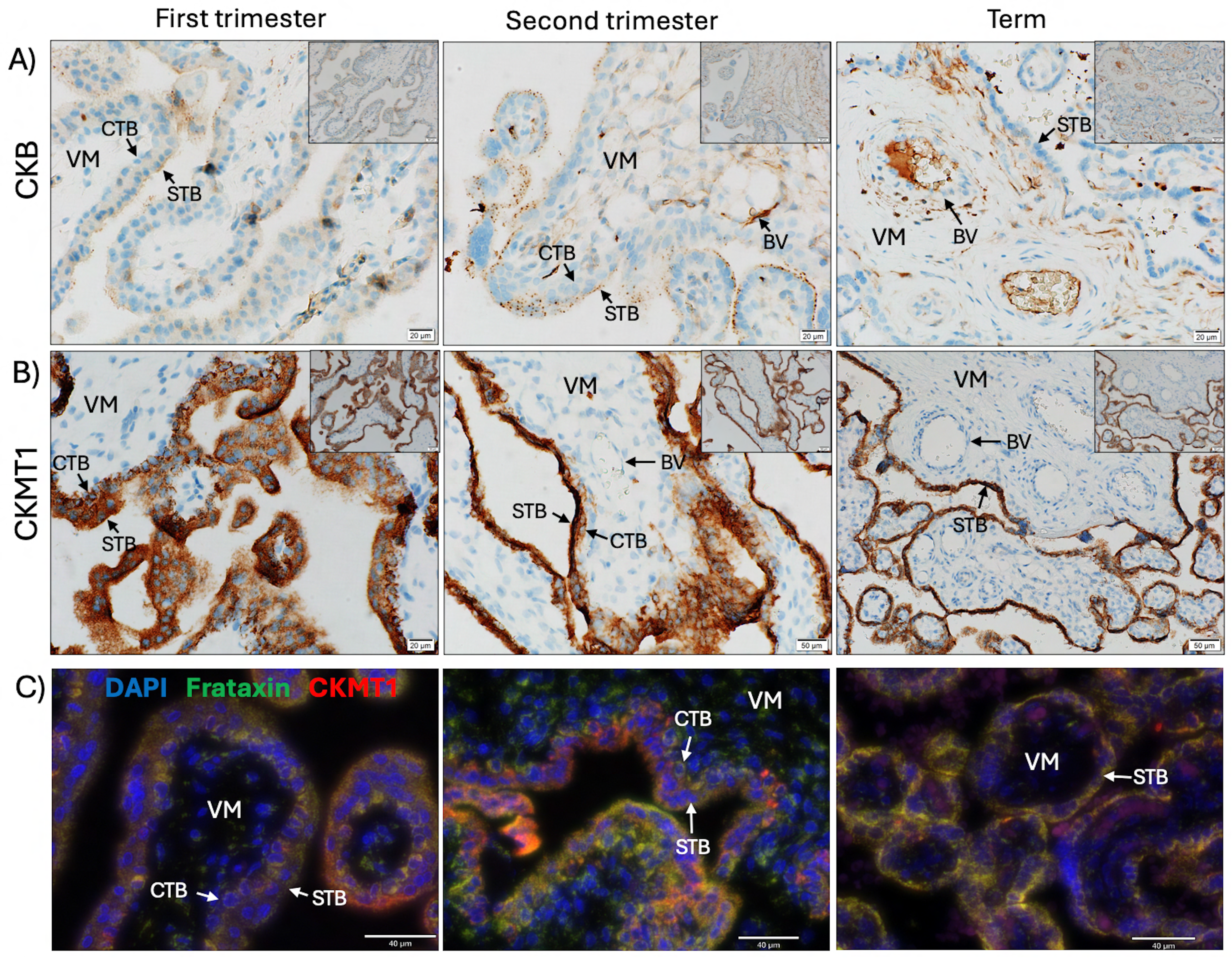
Immunolocalization of proteins associated with utilization of creatine or regeneration of ATP in human placenta across gestation. A) CKB localized to villous CTB and STB in first trimester, to the villous trophoblast as well as villous mesenchyme (VM) in second trimester, and to the VM including blood vessels (BV) and stromal cells in term placentae. B) CKMT1 expression was exclusive to the villous trophoblasts including the CTB and STB, without any detectable staining in the VM. C) Dual immunofluorescence showing co-localization of CKMT1 with the mitochondrial marker Frataxin across gestation. Scale bar 40μm. Main images of IHC were taken at 40x (scale bar 20μm), and the insets were images at 20x (scale bar 50µm). Images are representative of first trimester (n=6), second trimester (n=3) and term (n=6) placentae.

Immunofluorescence co-staining of CKMT1 with Frataxin (FXN), confirmed the mitochondrial localization of CKMT1 protein in the placenta (Fig 2C).

### Expression of the components of the creatine-CK-phosphocreatine system in primary trophoblast stem cells

Primary hTSCs are a well-established *in vitro* model to study mechanisms of placental development [32,33]. We first verified that primary hTSCs express creatine kinase, the central enzyme in the creatine phosphagen system. CKMT1 was expressed in primary hTSCs in a punctate pattern suggestive of mitochondrial localization (Fig 3A). Interestingly, it was localized in proliferating hTSCs expressing Ki67 as well as some spontaneously differentiating hTSCs that were negative for Ki67 (Fig 3B). This indicates that creatine phosphagen system may be important for both trophoblast proliferation and differentiation.

**Figure 3:**
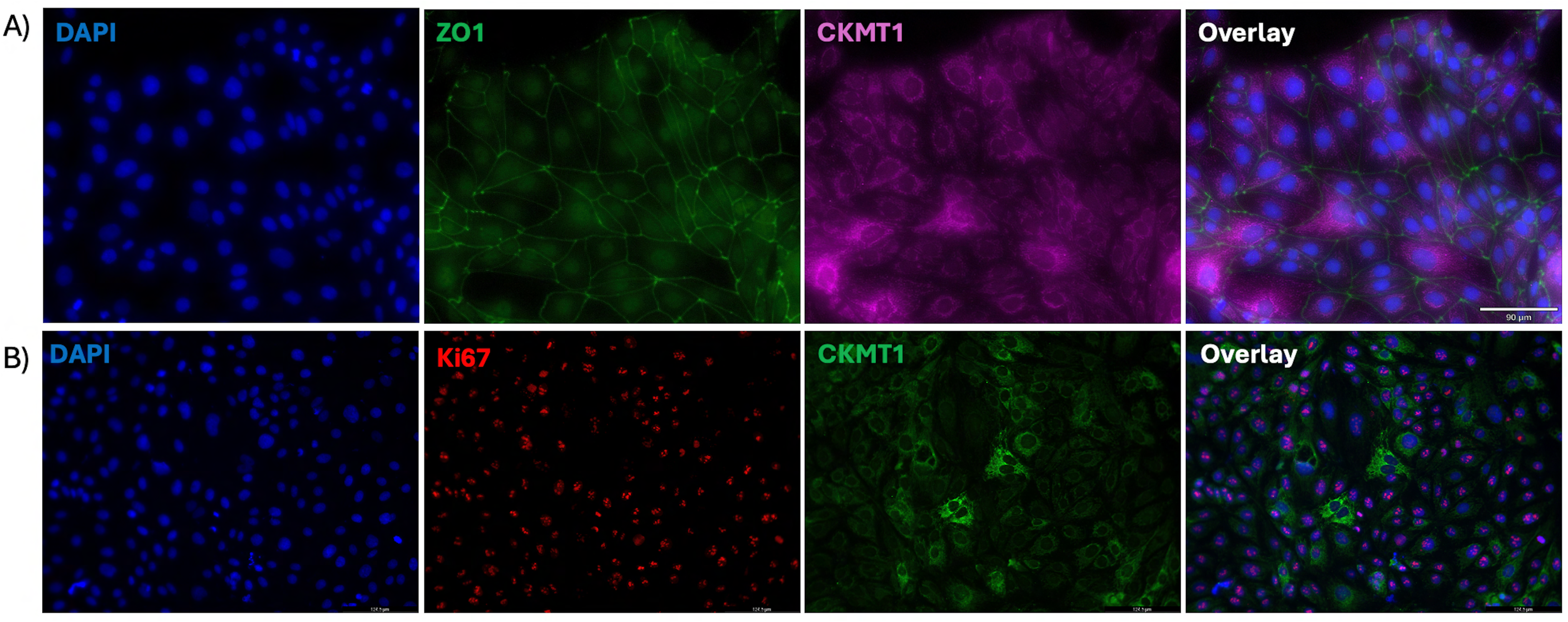
Immunolocalization of mitochondrial creatine kinase (CKMT1) in primary human trophoblast stem cells. A) Expression of CKMT1 with punctate staining indicative of mitochondrial localization in hTSC. Zona occludens-1 (ZO1) antibody used as a marker of cell membrane. Scale bar 90μm. Images are representative of 3 replicates of hTSC staining. B) Expression of CKMT1 in both proliferative hTSCs, marked with Ki67 protein, as well as spontaneously differentiating hTSCs (negative for Ki67). Scale bar 124.5µm. Images are representative of multiple images of 1 replicate of hTSC staining.

### Creatine-phosphocreatine metabolism influences trophoblast growth and phenotype

In this study, we observed that in the first trimester of pregnancy, the placental cytotrophoblasts express both the cytosolic and mitochondrial isoforms of creatine kinase. However, the exact role of the creatine kinase in placental development or function remains unknown. In our study, we modeled disruption of the Cr-CK-PCr system in primary hTSCs using the creatine analog and metabolic inhibitor of the creatine phosphagen system, cyclocreatine (cCr). Results shown in the main figures were generated using the 906p hTSC line, and these findings were further validated in 1000p hTSCs, with the corresponding data provided in the supplemental figures. Based on concentrations typically used in literature, we first tested the sensitivity of hTSCs to concentrations of cCr between 0.1 mM and 40 mM. We found a dose-response decrease in cell number after 48h of treatment compared to control (Fig. 4A and Supplemental Fig S4 A). Both cell lines showed no changes in cell number below 1mM and significant changes at concentrations above 10mM. Treatment of 906P hTSCs with 1 mM cCr did not impact cell count (P>0.05), but 10 and 20mM cCr conditions demonstrated a significant (*P* < 0.05) negative impact on total cell number at 48h of treatment (Fig 4B) compared to the 0mM cCr treatment (from hereon, referred as “control”). Treatment of the 1000p hTSC line also showed similar negative impact on trophoblast cell growth (Supplemental Fig S4 B and C). Since 20 mM cCr showed more robust and consistent results in both cell lines, most of further experiments, unless specified otherwise, were performed at this concentration.

**Figure 4:**
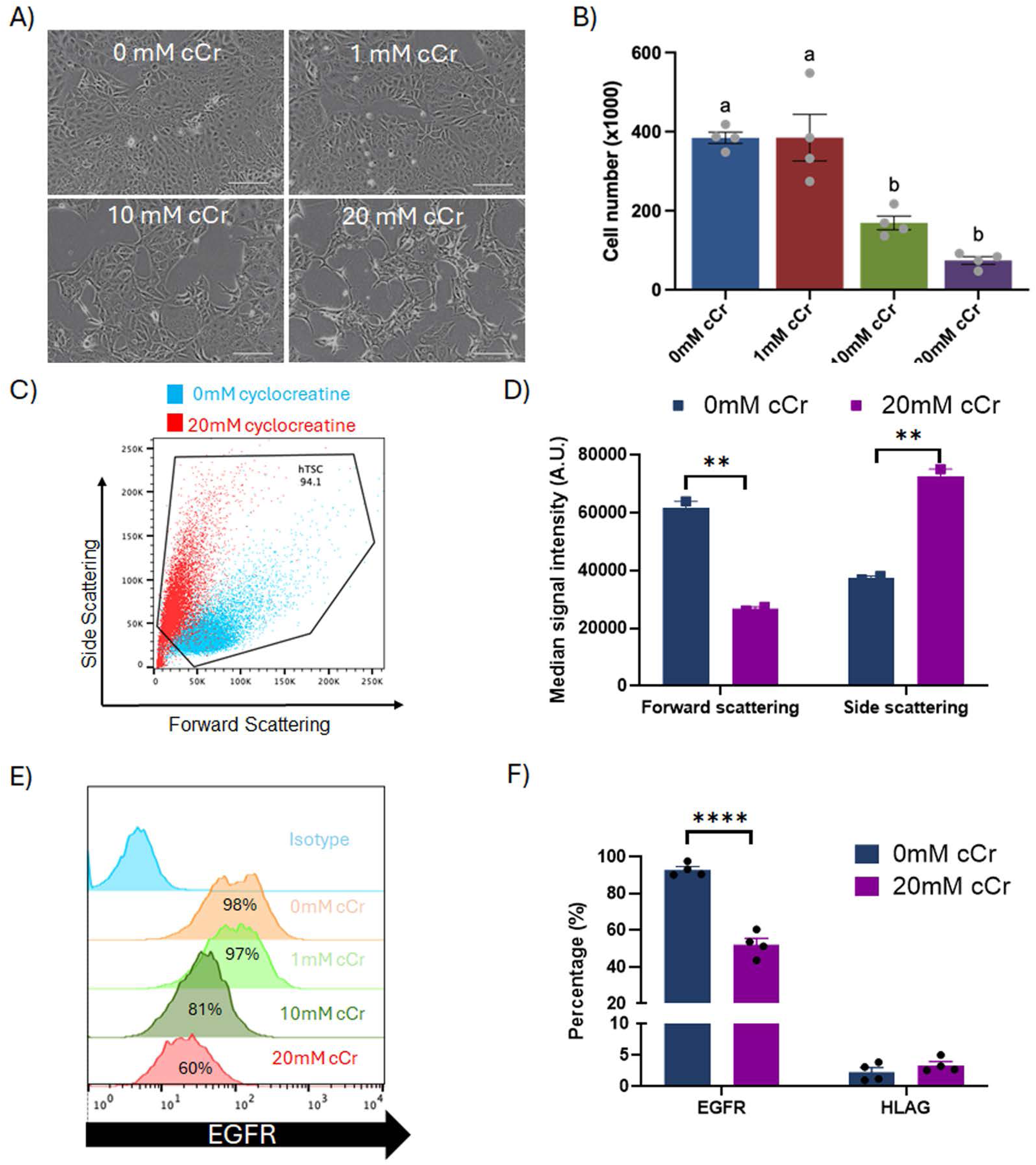
Cyclocreatine treatment alters hTSC morphology. A) Morphology of 906P hTSCs 48 h of culture in TSC complete containing 0, 1, 10, and 20 mM of cyclocreatine (cCr). Scale bar 160μm. B) 906P cell counts after treatment with different concentrations of cyclocreatine represented as mean ± SEM of 4 replicates. C) Flow cytometry analysis of 906P hTSCs in single cell suspension showing altered cell size (FSC) and granularity (SSC) following treatment with 20mM of cyclocreatine. D) Quantification of the FSC and SSC flow cytometry measurements of 906P hTSCs showing decreased forward-scattering and increased side-scattering following treatment with 20mM of cyclocreatine. E) Flow cytometry histograms showing percentages of 906P hTSC expressing cell-surface marker EGFR after treatment with different concentrations of cyclocreatine. F) Quantification of 906P hTSCs expressing EGFR and HLAG proteins by flow cytometry showing reduced proportions of EGFR positive hTSCs following treatment with 20mM of cyclocreatine. Bars represent median ± SEM of three replicates. Different letterheads on bars represent statistical significance. ** p≤0.01; **** p≤0.0001.

Cyclocreatine-treated cells appeared to have lost their epithelial phenotype, acquiring a more elongated shape (Fig 4A and S4B); therefore, we performed flow cytometry for the trophoblast epithelial cell marker, EGFR, and HLAG, a marker of EVT differentiation. During flow cytometry analysis, we noticed distinct cell clusters for the control vs cCr-treated hTSCs on the forward/side scatter plot, with a decline (P<0.01) in forward-scattering signal and a concurrent increase (P<0.01) in side-scattering signal in 20mM cCr-treated compared to control cells (Fig 4C and D). The 1000p hTSC line also revealed a decrease (P<0.001) in forward-scattering signal and an increase (P<0.001) in side-scattering signal in 20mM cCr-treated compared to control (Supplemental Fig S4D and E). Furthermore, while the proportion of cells expressing the EGFR protein was significantly decreased (P<0.0001) following cyclocreatine treatment (Fig 4E and F), there was no concurrent increase in HLA-G positive cells (< 5%) ruling out potential differentiation towards EVT. Similar results were obtained with 1000p hTSC line (Supplemental Fig 4F and G). The enzymatic activity of CK per cell in the 20mM cCr treated hTSCs was slightly greater (P<0.05) compared to the control (Supplemental Fig 4H) likely in response to the availability of additional substrate (cCr, a creatine analogue). These observations indicate an impairment of trophoblast growth and morphology following disruption of the creatine phosphagen system.

### Creatine-phosphocreatine metabolism affects trophoblast cell-cycle and apoptosis

Since there was an almost 3-fold decline in total cell number after cyclocreatine treatment, we investigated if the decline was due to interference in cell cycle or induction of apoptosis. First, a cell-cycle assay was done on hTSCs after 48h of culture in cCr medium. Flow cytometry analysis of EdU and DAPI fluorescence showed about 55% of the control hTSCs were in the S-phase (Fig 5A and B). In contrast, only about 7% of the hTSCs cultured in 20mM cCr medium were in the S-phase (P< 0.0001), while most of the cells were arrested in the either the G0/G1 (P< 0.0001) or G2/M (P< 0.0001) phase of the cell cycle (Fig 5A and B). A similar decline in cells in the S-phase with concurrent increase in cell arrest in the G0/G1 and G2/M phases were also seen in the 1000p hTSC line treated with 20mM cCr (Supplemental Fig 4I and J). To determine if disruption of the Cr-CK-PCr system triggers programmed cell death, we performed flow cytometry analysis of apoptosis using annexin V and PI staining of cyclocreatine-treated hTSCs. We observed a significant decline (P< 0.0001) in proportion of live cells with a concurrent increase (P< 0.0001) in apoptotic cells (Fig 5C and D) in the 20mM cCr-treated hTSCs compared to the control. In fact, continuous treatment of hTSCs with 20 mM cCr caused a siginficant decrease in cumulative cell number over 8 days (treatment-by-time interaction P=000.1) with increased dead cells floating in culture (Fig 5E). Therefore, interference in the creatine phosphagen system likely triggers cell cycle arrest as well as apoptosis resulting in reduced cell proliferation.

**Figure 5:**
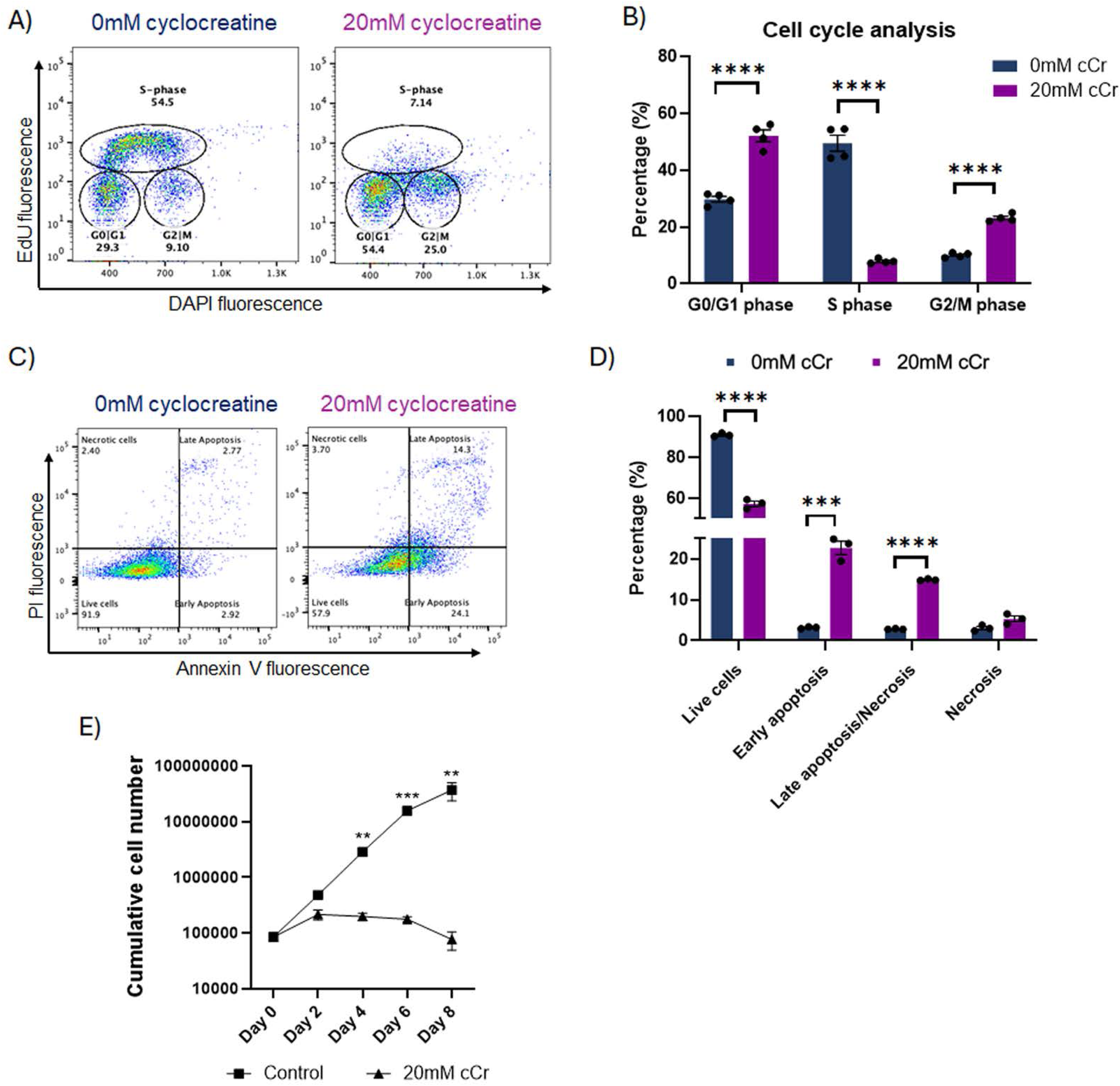
Cyclocreatine treatment causes hTSC cell cycle arrest and apoptosis. A and B) Flow cytometry analysis of cell-cycle in hTSCs showing a significant decline in proportion of cells in the S-phase with a concurrent increase in cells in the G0/G1 and G2/M phases of the cell cycle following treatment with 20mM of cyclocreatine. Bars represent mean ± SEM of four replicates. ****P ≤ 0.01. C and D). Flow cytometry analysis of cell apoptosis in hTSCs showing a significant decline in proportion of live cells with a concurrent increase in proportion of cells in the early and late apoptosis phase following treatment with 20mM of cyclocreatine. Bars represent mean ± SEM of three replicates. ***p ≤ 0.001; ****p≤0.0001 E) Cell count over 8 days of 20mM cyclocreatine treatment. Points represent mean ± SEM of three replicates. Two-way repeated-measures ANOVA **adj. p<0.01, ***adj p<0.001. Cyclocreatine significantly impaired cumulative hTSC growth over time, with a significant treatment-by-time interaction (p=0.0001).

### Creatine-phosphocreatine metabolism controls both oxidative phosphorylation and glycolytic energy metabolism in trophoblast stem cells

Placental trophoblasts can use oxidative phosphorylation as well as glycolysis for energy production [3]. To explore whether creatine kinase regulates energy metabolism of trophoblast cells, Seahorse mito-stress test was conducted following 48h of culture in cyclocreatine medium. With the cyclocreatine treatment, there was a dose-dependent decline (*P* < 0.01) in consumption of oxygen linked to basal respiration, maximal respiration, ATP production as well as coupling efficiency (Fig 6A and Supplemental Fig 5A). Additionally, a glycolytic stress test was performed to determine if there was any shift in energy metabolic pathway. The extracellular acidification rate linked with glycolytic rate (P<0.01), glycolytic capacity (P<0.001), and glycolytic reserve (P<0.001) were sharply diminished following cyclocreatine treatment (Fig 6B). Interestingly, the non-glycolytic acidification had the reverse trend with significantly higher non-glycolytic acidification (P<0.001) in cyclocreatine treated hTSCs compared to the controls (Fig 6B). The decline (P<0.001) in glycolysis related energy metabolism was also observed in the 1000p hTSC line (Supplemental Fig 5B). Taken together, these results indicate impairment of both OXPHOS and glycolytic pathways of energy production in hTSCs following inhibition of creatine phosphagen system. Interestingly, however, concentration of ATP on a per-cell level in the 20mM cCr-treated hTSCs was greater (P<0.05) than that in the control (Fig 6C), suggesting decline in ATP utilization/turn over in the dormant cells following cyclocreatine treatment.

**Figure 6:**
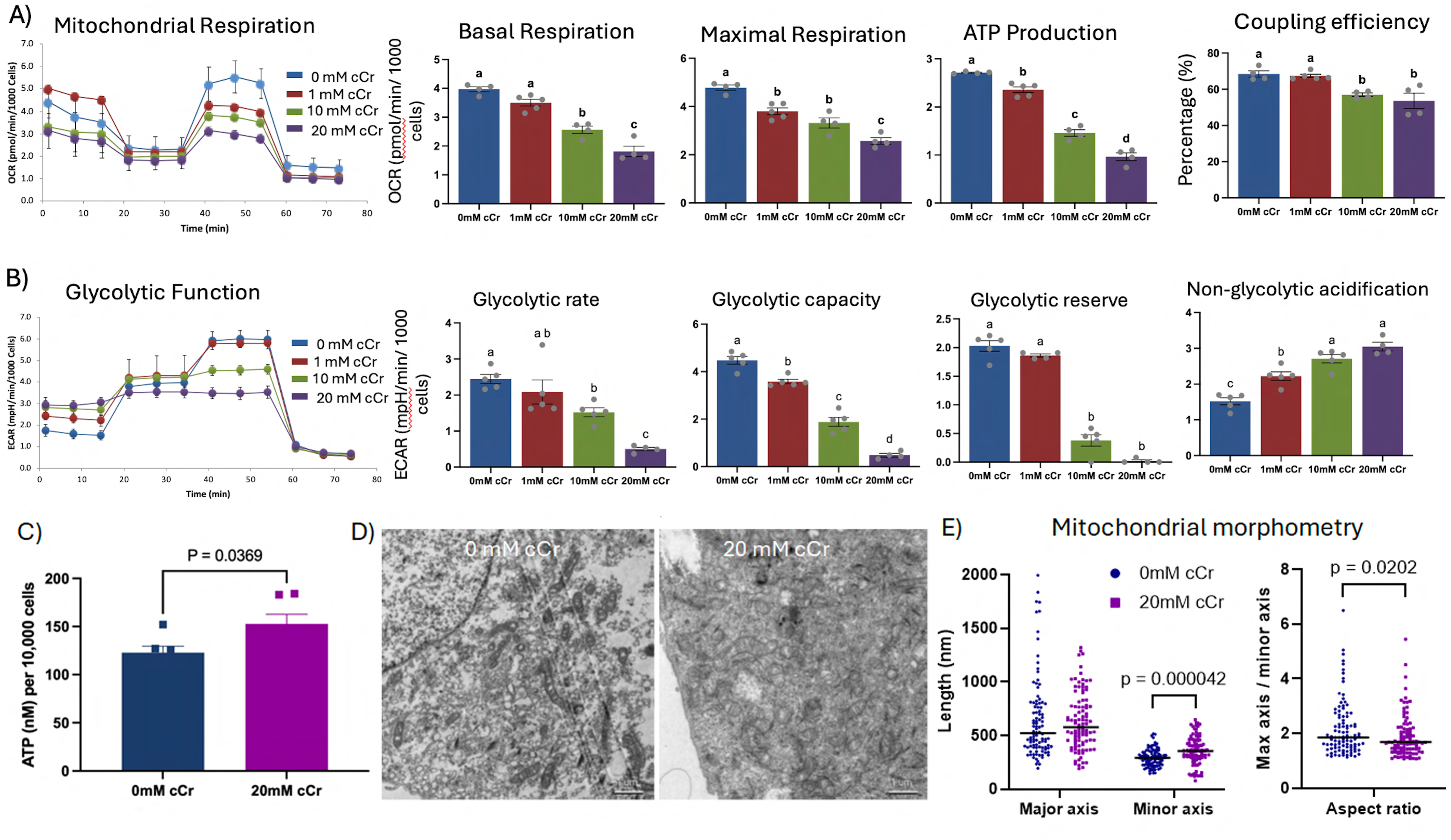
Cyclocreatine alters energy metabolism in hTSCs. A) Line graph and bar diagrams of Seahorse cell mito stress test assay in hTSCs following 48h of treatment with cyclocreatine showing decline in oxygen consumption rate (OCR) associated with basal and maximal respiration, ATP production, and coupling efficiency. B) Line graph and bar diagrams of Seahorse cell glycolytic-stress test assay in hTSCs following 48h of treatment with cyclocreatine showing decline in extracellular acidification rate (ECAR) associated with glycolytic rate, glycolytic capacity, and glycolytic reserve, but an increase in non-glycolytic acidification. C) Intracellular concentrations of ATP normalized to total cell number showing greater ATP content in cyclocreatine-treated hTSCs compared to the control. Bars represent mean ± SEM of 4-5 replicates. Different letterheads on bars represent statistical significance. D) Transmission electron microscopy images. E) Morphometry analysis of mitochondria from TEM images (n=100 mitochondria per treatment).

To further investigate mitochondrial health following 20mM cyclocreatine treatment for 48h, we performed transmission electron microscopy (TEM). Compared to controls cells, treated cells showed a slightly faint and swelled mitochondria (Fig 6 D-E), suggesting the beginning of compromised mitochondrial function.

### Influence of creatine-phosphocreatine metabolism disruption of gene expression in trophoblast stem cells

To explore potential mechanisms or pathways dysregulated by disruption of creatine phosphagen system, expression of genes involved in creatine metabolism, cell proliferation, apoptosis, redox potential, and cellular stress response were analyzed following 48h of cyclocreatine treatment (Supplemental Figure S6). Culturing the hTSCs in cyclocreatine medium did not influence (P>0.05) expression of genes associated with biosynthesis (*AGAT* and *GAMT*) or utilization (*CKB* and *CKMT1*) of creatine; however, a decline in *SLC6A8* gene expression was seen with 1mM (P<0.05) and 20mM (P<0.01) cCr treatment (Supplemental Figure S6A). Though the cell proliferation marker gene *MKI67* remain unchanged, expression of *PCNA* was significantly downregulated (P<0.05) in 20mM cCr condition (Supplemental Figure S6B). Expression of apoptotic marker genes *CASP3*, *CASP9*, and the *BAX/BCL2* gene ratio in hTSCs remained unchanged (P>0.05) with 10 and 20mM cCr treatment (Supplemental Figure S5B). Genes associated with redox capacity including SOD1 and GPX1 remained stable (P>0.05), however, expression of GPX8 declined significantly (P<0.05) with 20mM cCr treatment compared to the control (Supplemental Figure S6B). Expression of genes involved in cellular stress such as *MAPK8*, *MAPK9*, *P53*, *AMPK* remained unchanged (P>0.05) with cyclocreatine treatment (Supplemental Figure S6C). Similar genes associated with ATP synthesis (*ATP5F1B*), and mitochondrial dynamics (*MFN2* and *FIS1*) remained stable (P>0.05) in hTSCs following cyclocreatine exposure (Supplemental Figure S6C). These gene expression data demonstrate minimal influence of creatine kinase on transcriptional activity in hTSCs. However, reduced expression of *SLC6A8*, *PCNA*, and *GPX8* likely suggests a cellular and metabolic response of hTSCs to the sub-optimal function of creatine kinase in those cells.

### Creatine-phosphocreatine metabolism controls growth of primary trophoblast organoids

Primary human trophoblast organoids (TO) recapitulate 3D villous formation and aid in the *in vitro* assessment of physiological and developmental process of placental formation [34]. We used primary TO derived from first trimester placental tissue to confirm the role of the creatine phosphagen system in placental development. TOs cultured in cyclocreatine medium showed delayed growth with visibly smaller organoids, relative to control TOs, by day 6 of treatment (Fig 7A). Cell count of the dissociated TO in single cell suspension demonstrated significant reductions (P<0.05) in cell numbers in the cCr-treated TO compared to the controls (Fig 7B). The proportions of cells in the TO expressing EGFR were also lower (P<0.05) in the cCr treated compared to the control organoids (Fig 7C).

**Figure 7:**
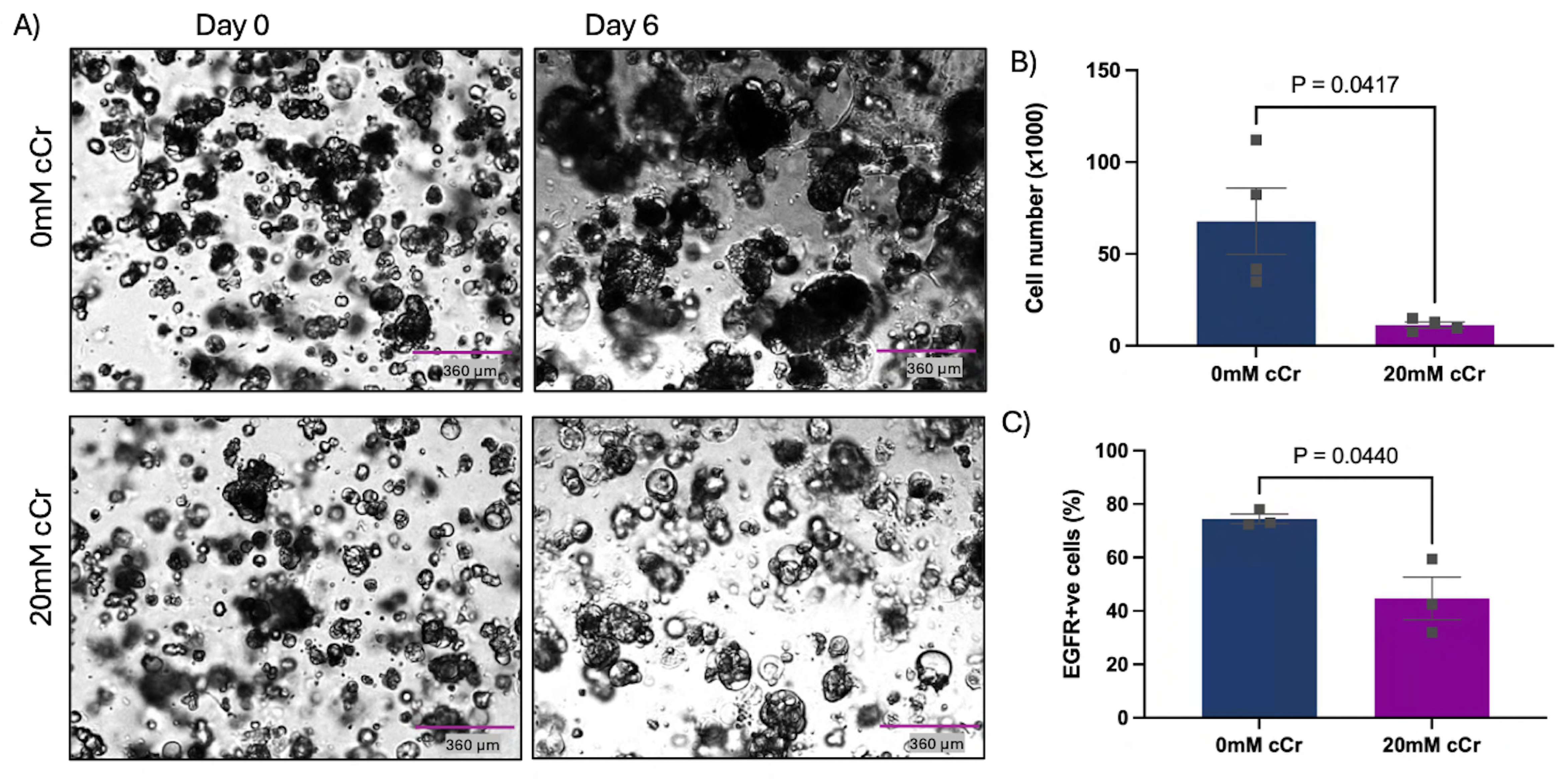
Cyclocreatine treatment affects primary human trophoblast organoid (TO) growth. A) Growth of TO following six days of culture in iCTB complete containing 0 or 20 mM of cyclocreatine (cCr) showing reduced organoid formation and growth with cCr treatment. B) Cell counts in the single-cell suspension of TO was reduced following cCr treatment. C) EGFR expression in TO cells was reduced following cCr treatment. Bars represent mean ± SEM of 3-4 replicates.

### Creatine supplementation partially restores growth and epithelial phenotype of cyclocreatine treated primary trophoblast stem cells

Cyclocreatine is a creatine analogue and a competitive inhibitor of cellular creatine uptake [20]. Therefore, we used creatine, at varying concentrations, alongside the cyclocreatine treatment to investigate if creatine can negate the detrimental effects of cyclocreatine on hTSC growth and phenotype. We first conducted an experiment with 20mM cyclocreatine and increasing concentrations (1, 5, 10, 20mM) of creatine; however, the combinations proved to be more detrimental (data not shown) potentially due to transporter (SLC6A8) saturation, osmolarity changes, or a combination thereof. The negative impacts of CK inhibition on trophoblast growth were also observed with 10mM cCr treatment so, we repeated the experiment with 10mM cCr (Fig 8). As expected, the total cell number decreased with 10mM cCr treatment compared to the control (0mM cCr). Supplementation with creatine, alongside the presence of cyclocreatine, improved cell growth and elevated the total cell number with significant difference (P<0.05) at 1mM creatine compared to 10mM cCr alone (Fig 8A and B). Cyclocreatine treatment resulted in hTSCs with lower forward-scattering (P<0.01) and greater side-scattering (P<0.01) of cells, whereas the addition of creatine at 5mM and greater concentrations to the cyclocreatine medium reverted back (P<0.01) this altered phenotype similar to that of control hTSCs (Fig 8C, E and F). The proportion of cells expressing EGFR declined (P<0.01) with 10mM cCr treatment (Fig 8D and G) as anticipated. The addition of creatine (at all tested concentrations) elevated (P<0.05) the composition of EGFR expressing cells (Fig 8D and G) although it remained lower (P< 0.05) compared to the control hTSCs (Fig 8H). There was a slight increase (P< 0.05) in the percentage of cells expression HLA-G following the supplementation of creatine; however, those were still less than 5% of the total cell population (Supplemental Figure S6D). Overall, these data suggest a role for the creatine phosphagen system in supporting trophoblast cell function in the placenta across gestation (Fig 9).

**Figure 8:**
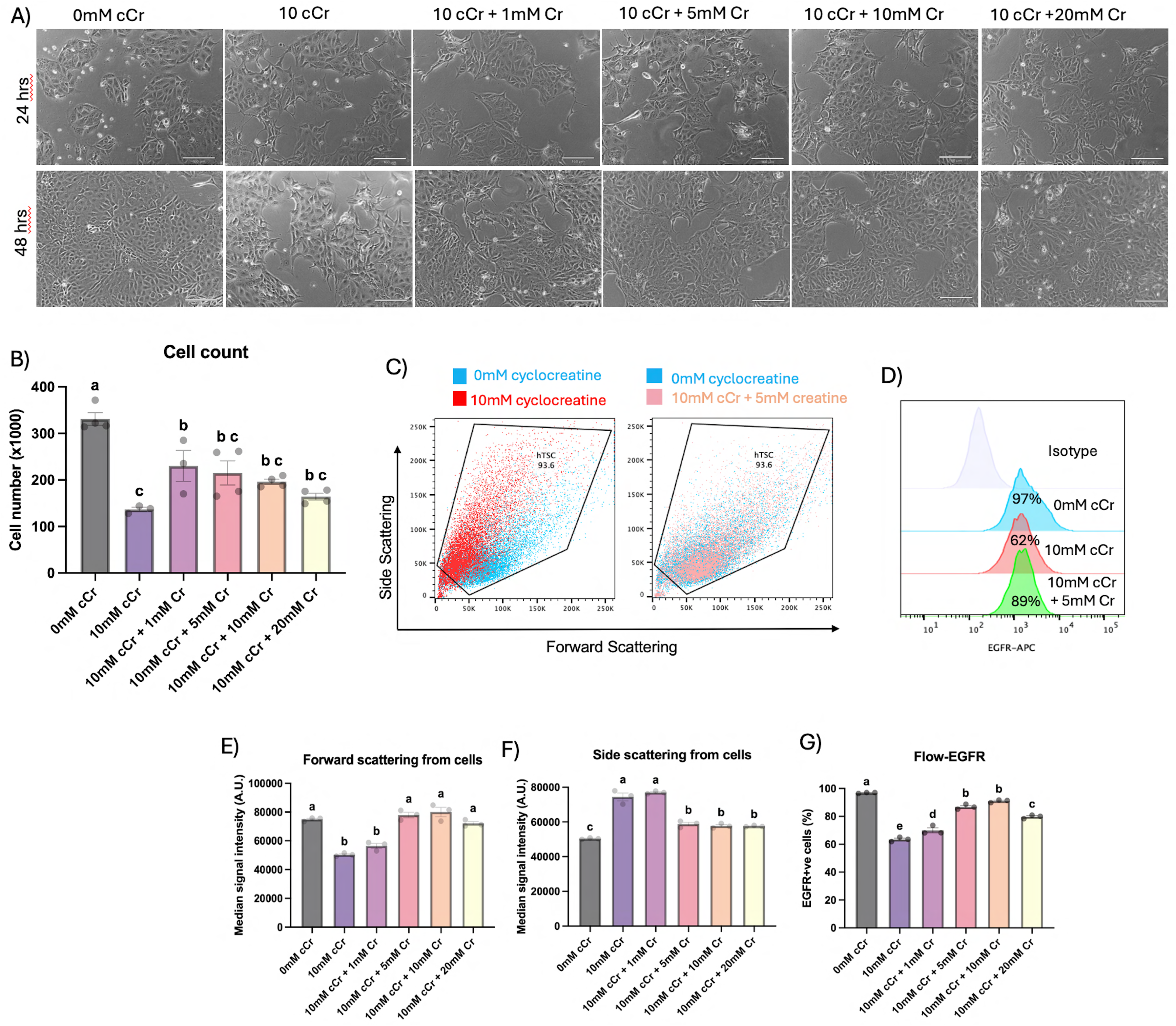
Creatine addition prevents cyclocreatine effects on hTSCs. A) Cell morphology of hTSCs treated with 10mM cyclocreatine along with or without creatine at various concentrations. Scale bar 160μm B) Cell counts following 48h of cyclocreatine treatment with/without creatine showing a decline in cell number with 10mM cyclocreatine but an increase in cell number with the addition of 1mM creatine. Bars represent mean ± SEM of 3-4 replicates. C) Flow cytometry analysis of hTSCs showing altered cell size (FSC) and granularity (SSC) following treatment with 10mM of cyclocreatine whereas the simultaneous addition of 5mM creatine reverse the cell phenotype to normal range (0mM cyclocreatine). D) Flow cytometry histograms showing percentages of hTSCs expressing EGFR cell-surface marker in hTSCs after 10mM cyclocreatine treatment along with of 5mM creatine. E and F) Quantification of the forward and side-scattering of flow cytometry measurements of hTSCs showing decreased forward-scattering and increased side-scattering following treatment with 10mM of cyclocreatine. After the addition of 5mM creatine, the cell size and granularity are reversed to normal levels as comparable to 0mM cCr. Bars represent median ± SEM of three replicates. Different letterheads on bars represent statistical significance. G) Quantification of hTSCs expressing the cell-surface marker EGFR by flow cytometry showing reduced proportions of EGFR-positive hTSCs following treatment with 10mM of cyclocreatine; however, with the addition of 5mM creatine, the proportions of EGFR positive cells increase significant but still falls below normal levels (0mM cCr). Bars represent mean ± SEM of three replicates. Different letterheads on bars represent statistical significance.

**Figure 9:**
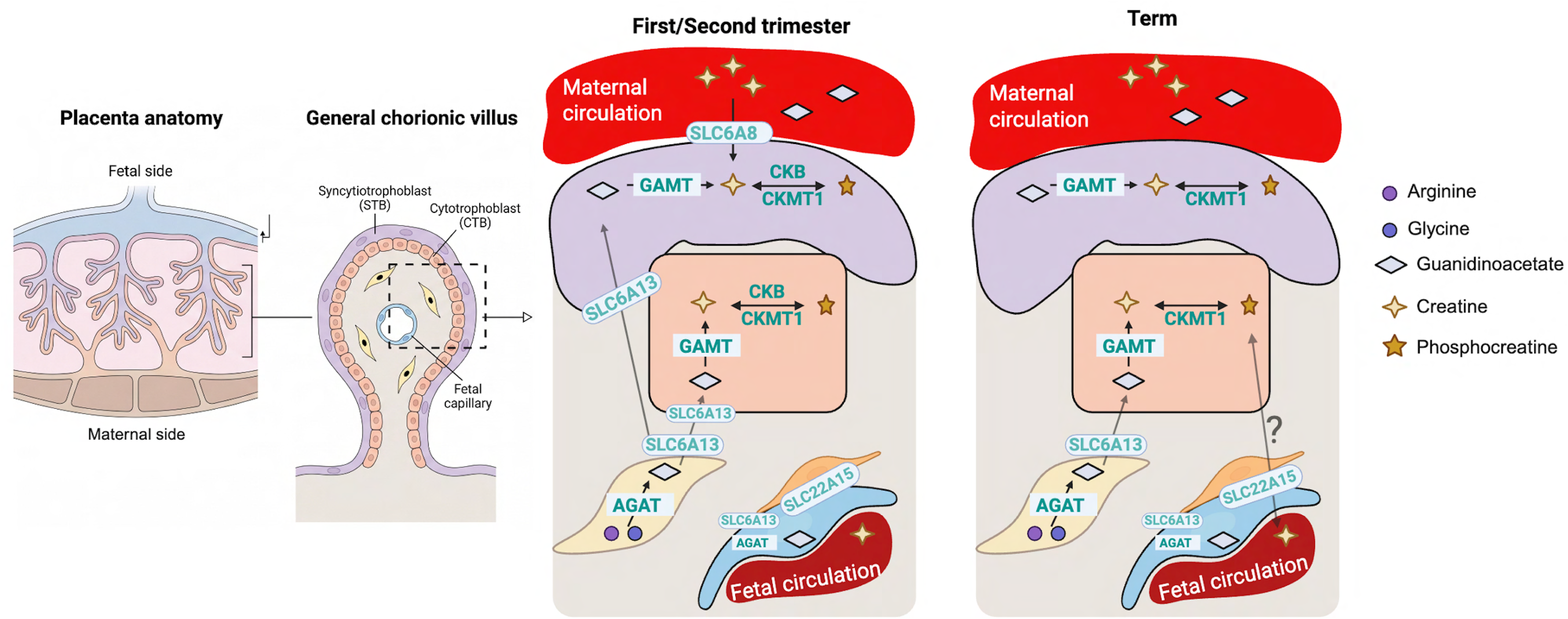
Graphical Abstract. (Figure created in BioRender.com/q7vmeqb).

## Discussion

The creatine phosphagen system is known to regulate ATP homeostasis in tissues with high energy demand such as skeletal muscle and brain [35]. The placenta is a tissue with high metabolic and bioenergetic demand owing to its development process such as cell proliferation, fusion, and invasion as well as functional needs such as macromolecule biosynthesis, and nutrient secretion and transport [2,3]. The presence of creatine in human placenta was first reported by Mischel in 1955, who demonstrated measurable concentrations of both creatine and creatinine (a metabolite of creatine) in placental tissue [36]. Years later, multiple studies confirmed the presence of different components of the creatine phosphagen system in human placenta [10,24,25,37–40]. But it was not until 2017 that Ellery and colleagues demonstrated the cellular localization of the proteins for creatine biosynthesis and transport using term human placenta tissues [10]. To date, the knowledge of the different components (biosynthetic, transport, and utilization) of this system at cellular level remains unclear, especially earlier in gestation when placental development is rapid. This study is the first to provide a complete cellular picture of the biosynthesis, transport and utilization of creatine by human placenta from early first trimester to term (Fig 9). Additionally, our study is also the first to provide direct evidence for the role of the creatine phosphagen system in placental trophoblast cell growth.

Tissues with high ATP demand such as brain [6], testes [8], and gut [9] are known to synthesize creatine locally. The placenta at term is also capable of synthesizing creatine, predominantly by the trophoblast cells [10]. We found that the human placenta, throughout gestation, may be capable of metabolizing creatine. The biosynthesis, transport, or utilization of creatine did not appear to be influenced by fetal sex in term placenta in this study, as was noted previously [10]. Similarly, no differences were noted in early vs late first trimester placentae for the different components of the creatine phosphagen system. In line with previous report [10], we observed that, in the term placenta, the protein responsible for direct creatine synthesis, i.e. GAMT, was expressed mostly by the STB, whereas the protein for synthesis of the creatine precursor-guanidinoacetate (GA), i.e. AGAT, was localized only in the villous stromal cells and endothelial cells of fetal capillaries but not expressed by villous trophoblast cells. We saw similar staining pattern in placentae from first and second trimester with additional confirmation of GAMT expression by the CTB layer. This indicates a co-operation between the villous mesenchymal and trophoblast compartments for the local biosynthesis of creatine in placenta throughout gestation. This concept is further supported by the histochemical localization of SLC6A13, a transporter capable of importing GA [41], in the villous CTBs and STBs, specifically, in the first and second trimester placentae observed in this study indicating a possible shuttling of creatine precursor and products between placental regions.

Ellery and colleagues showed the localization of the creatine transporter, SLC6A8, in villous STB layer in term placenta. Our study does not support this previous finding wherein the differences may be due to variability in antibody specificity, epitope recognition, or experimental conditions used between studies. Our study, though, revealed the expression of SLC6A8 at the apical surfaces (likely to microvilli) of STB in first trimester placenta suggesting that the trophoblast cells of the placenta may depend on maternal creatine early in gestation and then become self-reliant for creatine synthesis later in gestation. Moreover, the localization of SLC22A15, a newly identified exporter of creatine [5], in the pericytes and endothelial cells lining the fetal blood vessels may support the export of creatine from placental/fetal compartment, the directionality of which requires further experimental confirmation (Fig 9). The placentae at all three trimesters expressed the creatine kinases in villous CTB and STB indicating that they may also use the creatine phosphagen system for ATP recycling. Overall, this study builds new knowledge about the creatine phosphagen system in placenta across gestation suggesting its importance in placental bioenergetics and development.

In this study, we further evaluated the role of the Cr-Ck-PCr system in trophoblast growth using first trimester hTSCs. Cyclocreatine is a creatine analogue that competes with cellular creatine uptake and inhibits the function of creatine phosphagen system in multiple model systems [14,15,17,19,20,42–44]. Normally, cells are in equilibrium with Cr and PCr at 1:1 ratio, but cyclocreatine competes with creatine both for cellular uptake and as a substrate for CK, rapidly forming cyclocreatine phosphate (cCrP) at a much higher ratio (10:1 for cCr and cCrP) but with lower affinity to regenerate ATP, and therefore act as a sink for ATP [14,15]. Therefore, cyclocreatine interferes with the normal Cr-CK-PCr pathway, but specifically with PCr metabolism [20,43]. We observed that treatment of hTSCs with 10 or 20 mM cyclocreatine significantly impaired cell growth, primarily through reduced proliferation. Disruption of creatine metabolism may limit ATP availability and/or buffering capacity, thereby imposing metabolic constraints on cell division. Interestingly, in glioblastoma, PCr was recently shown to promote proliferation through an epigenetic mechanism involving stabilization of Bromodomain-Containing Protein 2 (BRD2), a regulator of chromosome segregation[43]. Whether this PCr– BRD2 epigenetic mechanism similarly contributes to trophoblast proliferation remains to be investigated. The morphology of hTSCs treated with cyclocreatine was also altered wherein the cells become more elongated and less epithelial-like phenotype. This observation was further corroborated by a significant decrease in their cell size (measure of FSC) and increased internal complexity or granularity (measure of SSC) on flow cytometry analysis. This pattern suggests that perturbation of creatine-dependent energy homeostasis induced a stress-associated cellular remodeling phenotype, potentially reflecting cell shrinkage, cytoplasmic condensation, altered organelle structure, growth arrest and/or early apoptotic changes. Indeed, recent study by Chen and colleagues (2026) using embryonic stem cells demonstrated that inhibition of Cr-PCr metabolism by cyclocreatine disrupts normal cellular cytoskeletal organization, via “energy buffering-F actin assembly-YAP nuclear localization” axis leading to developmental arrest of blastocysts [44]. Alteration of cellular phenotype has also been observed in pancreatic adenocarcinoma cells treated with cyclocreatine [18]. In our system, the loss of trophoblast epithelial morphology was further confirmed by the significant reduction in trophoblast epithelial marker EGFR expression in cyclocreatine-treated hTSCs. Similar results were observed using trophoblast organoids, suggesting that this effect is specific to stem cells/progenitors of the placenta. In literature, tumor cell spheroids treated with cyclocreatine showed decline in cell invasion, a process requiring cytoskeleton reorganization and ATP, similar to that required for cell division [18]. Collectively, these results suggest that creatine-dependent phosphate buffering contributes to maintaining the energetic environment necessary for trophoblast progenitor activity and tissue growth.

We also determined the effect of cyclocreatine on CK activity and, interestingly, we found that the CK activity per cell in hTSCs was slightly increased with 20mM cyclocreatine treatment, likely due to greater rate of forward reaction of CK in the presence of higher concentrations of cellular cyclocreatine. Whether cyclocreatine or the new product, cyclocreatine phosphate [15], influences the rate of reverse reaction of CK remains unknown based on the working principle of the CK activity kit used in this study.

Since cyclocreatine impaired the normal proliferation of the hTSCs, we performed cell cycle and apoptosis assays to understand the cause of reduced cell growth. Cyclocreatine treatment significantly reduced the proportion of cells in S-phase while increasing the fraction of cells in G0/G1 and G2/M phases. This shift in cell-cycle distribution suggests impaired cell-cycle progression, with cells accumulating at the G1/S and G2/M checkpoints. Successful cell division requires rearrangement of actin cytoskeleton and correct segregation of duplicated chromosomes wherein any disorganization during the cytokinesis stage can lead to polyploidy [45]. Phosphocreatine is known to provide ATP for actin phosphorylation and rearrangement [46]. Given the role of the Cr-CK-PCr system in ATP recycling, our findings are consistent with energetic stress limiting DNA replication and mitotic progression, ultimately suppressing trophoblast proliferation. This finding is in agreement with multiple other studies using cancer cell models [19,20,42]. Cyclocreatine treatment of hTSCs also resulted in an increase in apoptotic cells, which has been observed previously in tumor cell lines [47,48]. The combination of reduced S-phase entry, accumulation of cells in G0/G1 and G2/M phases, and increased apoptosis is consistent with metabolic stress activating cell-cycle checkpoints, with a subset of cells subsequently undergoing programmed cell death likely due to failure of sustaining the prolonged energetic stress to maintain their basic membrane integrity and ion gradient [49].

Previous studies have shown that the cell cycle arrest following functional inhibition of CK is associated with a simultaneous decline in ATP metabolism [50,51]. In this study, Seahorse analysis demonstrated that cyclocreatine impairs trophoblast bioenergetics in a dose-dependent manner. cCr treatment progressively reduced basal respiration, maximal respiration, and ATP-linked oxidative phosphorylation, indicating compromised mitochondrial energy production and reduced respiratory reserve. In parallel, glycolytic rate, glycolytic capacity, and glycolytic reserve were also diminished, particularly at higher cCr concentrations, suggesting that trophoblast cells were unable to compensate for mitochondrial dysfunction through increased glycolytic flux. Although mitochondrial cristae morphology remained preserved following 48 h of cCr treatment, mitochondria exhibited reduced electron density and mild matrix swelling relative to control, consistent with early ultrastructural alterations indicative of early-stage mitochondrial stress.

Interestingly, we did not observe decreased ATP per cell, which was modestly increased instead. Previous studies in tumor cells have reported a similar phenomenon, in which cyclocreatine treatment impairs both mitochondrial and glycolytic ATP production [52] without substantial reduction in steady-state ATP levels. Instead, they observed decrease in PCr and increase in the ADP/ATP ratio leading to activation of AMPK signaling [18,53]. Cyclocreatine competes with creatine for phosphorylation by creatine kinase, resulting in the formation of cyclocreatine phosphate, which may function as a less efficient phosphagen compared with phosphocreatine [54] for the regeneration of ATP. Consequently, although ATP may be partially maintained, the buffering capacity of the creatine-phosphocreatine system is disrupted, leading to energetic stress that activates cellular energy-sensing pathways such as AMPK [53,55]. Our results are consistent with this model of disrupted high-energy phosphate buffering. The increase in ATP per cell despite reduced mitochondrial and glycolytic capacity likely reflects reduced ATP consumption associated with suppressed cell proliferation. In addition, Seahorse analysis revealed an increase in non-glycolytic acidification in cCr-treated cells, suggesting broader metabolic remodeling and activation of stress-associated metabolic processes rather than productive glycolytic ATP generation. Taken together, these findings indicate that cyclocreatine disrupts creatine-dependent energy buffering in trophoblast cells, leading to reduced metabolic flexibility and activation of energy stress responses. Under these conditions, surviving cells appear to maintain a minimal ATP pool through reduced ATP demand and limited phosphate transfer from cyclocreatine phosphate, while overall bioenergetic capacity remains impaired. Cyclocreatine is transported through a sodium-dependent active transport SLC6A8, but does not affect membrane integrity or cellular ATP content [53]. In cells with high-glucose dependency, the toxicity of cCr may be due to accumulation of cyclocreatine phosphate (cCrP) and cellular swelling [15,53], however this mechanism was not evident in other cell types. Recently, cyclocreatine has been shown to suppress phosphocreatine production, global protein-phosphorylation, and signal transduction [51,52]. Treatment of tumor cells with another CK inhibitor, dinitrofluorobenzene, has also been reported to suppress mitochondrial respiration, volume and membrane potential, and cell proliferation [51,52]. Based on previous studies and our current findings, it is highly plausible that the cyclocreatine blocks ATP homeostasis by the Cr-CK-PCr pathway leading to a broader decline in protein-phosphorylation for cellular growth and function, although further studies will be required to confirm this hypothesis. Indeed, supplementation of exogenous phosphocreatine or ATP restores normal cellular mitochondrial function, viability, and proliferation [50–52,56] further confirming the role of the creatine phosphagen system in ATP buffering and signal transduction.

We also determined if the growth inhibition caused by cyclocreatine resulted from disruption of the Cr-CK-PCr system rather than nonspecific toxicity. Any improvements following the exogenous supplementation of either creatine or phosphocreatine in the cyclocreatine treated cells would confirm the interference of the cCr with the creatine-phosphagen system and ATP recycling. Multiple studies have shown that supplementation of phosphocreatine indeed alleviates the negative impacts of cyclocreatine treatment [50,52,56,57]. However, the specific mechanism of transport of PCr into cells is still unknown. Therefore, in this study, we tested if exogenous creatine could negate the impacts of cyclocreatine and restore cellular proliferation and phenotype. Trophoblast cells were treated with 10mM cCr in the presence of increasing concentrations of creatine. Co-treatment with creatine partially rescued the inhibitory effects of cCr on trophoblast growth, as reflected by increased cell number compared with cCr treatment alone. Creatine supplementation also restored cellular morphology, as indicated by recovery of forward and side scatter profiles, suggesting normalization of cell size and internal complexity. Importantly, the proportion of EGFR-positive cells, which was markedly reduced by cCr treatment, increased substantially in the presence of creatine. These findings indicate that the negative effects of cyclocreatine on trophoblast proliferation and progenitor maintenance are largely attributable to competition with endogenous creatine and disruption of creatine-dependent phosphate buffering. Together with the observed reductions in mitochondrial respiration, glycolytic capacity, and trophoblast stem cells and organoid expansion, these results support a model in which the creatine phosphagen system plays an important role in maintaining trophoblast metabolic flexibility and proliferative capacity. Disruption of this pathway by cyclocreatine places trophoblast cells in an energetically constrained state that limits progenitor expansion and tissue growth. Disruption in the creatine phosphagen system has been linked to multiple pregnancy-associated disorders such as fetal growth restriction, preeclampsia, and gestational diabetes [3,24–27]. Therefore, our study provides fundamental knowledge to investigate the causal effect of the creatine phosphagen system and rescue strategies, perhaps with creatine supplementation, in cases of placental disorders.

## Supplementary data

All supplementary data relevant to this manuscript is available online at *Human Reproduction*.

## Authors’ contribution

The study design and experimental planning was done by N.S. and F.S. and executed by N.S with assistance from M.M. (Placental tissue collection), R.R. and D.P. (IHC and IF staining), W.B.S. (Seahorse assay), F.H.S. (Organoid experiment and TEM) and C.Z., M.B.N., and L.M.M. (cell maintenance and count). The first draft of the manuscript was written by N.S., F.H.S., D.P., and F.S. and edited by all the coauthors. The final version of the manuscript was prepared by N.S. and F.S. All authors read and approved the final manuscript.

## Data availability

All datasets generated and analyzed in this study are included in the published article.

## Acknowledgements

The authors thank the patients for their voluntary donation of placental tissues for research. The Seahorse assay analysis was performed at Mitochondrial Bioenergetics Resource Core at University of California San Diego. The authors would like to thank the University of California, San Diego - Cellular and Molecular Medicine Electron Microscopy Core (UCSD-CMM-EM Core, RRID:SCR_022039) for equipment access and technical assistance.

---

**Supplementary Figure 1:** No primary controls for immunohistochemistry in placenta tissues across gestation for anti-rabbit (A and B) and anti-mouse (C) antibodies.

**Supplementary Figure 2:** Immunofluorescence localization of proteins associated with synthesis and utilization of creatine in human placenta across gestation. B) GAMT expression was observed mainly in the placental villous cytotrophoblast (CTB) and syncytiotrophoblast (STB) across gestation. B) CKB localized mostly to apical surfaces of STB in first and second trimester but absent in STB from term placentae. Images of IF staining were taken at 20x (scale bar 60µm. Images are representative of first trimester (n=6), second trimester (n=3) and term (n=6) placentae.

**Supplementary Figure 3:** Immunolocalization of proteins associated with transport of creatine and its precursor in human placenta across gestation. A) SLC6A13 linked to transport of guanidinoacetate was localized to both villous CTB and STB and villous mesenchyme (VM) including blood vessels (BV) and stromal cells. B) SLC22A15, a newly reported transported linked to export of creatine was localized mainly to the endothelial cells lining the blood vessels (BV) across gestation, more prominently in the second trimester and term placentae. Main images of IHC were taken at 40x (scale bar 20μm), and the insets were images at 20x (scale bar 50μm). Images are representative of first trimester (n=6), second trimester (n=3) and term (n=6) placentae.

**Supplementary Figure 4:** Treatment of hTSCs with cyclocreatine affect cell proliferation and morphology in a dose-dependent manner. A) Cell number following a dose-response treatment of 906P and 1000P hTSCs with cyclocreatine concentrations between 0.1 mM ad 40mM. Data are shown as mean ± SEM of 8 replicates. Different letters mark statistically significantly different treatments. B) Morphology of 1000P hTSCs after 24 and 48 h of culture in TSC complete containing 0 and 20 mM of cyclocreatine (cCr). C) Cell counts after treatment of 1000P hTSCs with 0 or 20mM cyclocreatine represented as mean ± SEM of 4 replicates. D) Flow cytometry analysis of 1000P hTSCs showing altered cell size (FSC) and granularity (SSC) following treatment with 20mM of cyclocreatine. E) Quantification of the FSC and SSC flow cytometry measurements of 1000P hTSCs showing decreased forward-scattering and increased side-scattering following treatment with 0 or 20mM of cyclocreatine. Bars represent median ± SEM of three replicates. F) Flow cytometry histograms showing percentages of 1000P hTSCs expressing EGFR and HLA-G cell-surface markers after treatment with 0 or 20mM cyclocreatine. G) Quantification of 1000P hTSCs expressing EGFR and HLA-G proteins by flow cytometry showing reduced proportions of EGFR+ hTSCs following treatment with 20mM of cyclocreatine. Bars represent mean ± SEM of three replicates. H) Enzymatic activity of CK normalized to total cell number in control and 20mM cCr-treated 906P hTSCs. Bars represent mean ± SEM of four replicates. I-J) Flow cytometry analysis of cell cycle in 1000P hTSCs showing a decline in proportion of cells in the S-phase with a concurrent increase in cells in the G0/G1 phase of the cell cycle following treatment with 20mM of cyclocreatine. Bars represent mean ± SEM of two replicates.

**Supplemental Figure 5:** Validation of energy metabolism changes by cyclocreatine treatment in human trophoblast stem cell (hTSC) line 1000p. A) Line graph and bar diagrams of Seahorse cell mito stress test assay in hTSCs following 48h of treatment with 20mM cyclocreatine showing decline in oxygen consumption rate (OCR) associated with basal and maximal respiration, and ATP production. B) Line graph and bar diagrams of Seahorse cell glycolytic-stress test assay in hTSCs following 48h of treatment with 20mM cyclocreatine showing decline in extracellular acidification rate (ECAR) associated with glycolytic rate, glycolytic capacity, and glycolytic reserve. Bars represent mean ± SEM of 4-5 replicates. Different letterheads on bars represent statistical significance.

**Supplemental Figure 6:** Effects of cyclocreatine treatment on genes involved in creatine metabolism, cellular stress and antioxidant system in human trophoblast stem cell (hTSC) line 906p. A) Expression of genes associated with creatine metabolism showing a significant decline in creatine transporter *SLC6A8* with 20mM cyclocreatine treatment. B) Expression of genes associated with cell proliferation, apoptosis and redox potential showing a significant decline in *PCNA*, and *GPX8* but an increase in *CASP9* with 20mM cyclocreatine treatment compared to 0mM cyclocreatine. C) Expression of genes associated with cellular stress response showing unaltered transcriptional activity with 20mM cyclocreatine treatment. Bars represent mean ± SEM of 3-5 replicates. D) Quantification of hTSCs expressing the cell-surface marker HLA-G by flow cytometry showing unaltered HLAG+ hTSCs following treatment with 10mM of cyclocreatine; however, with the addition of 5mM creatine, the proportions of HLA-G+ cells increase significant but at a very low level compared to control (0mM cCr). Bars represent mean ± SEM of three replicates. Different letterheads on bars represent statistical significance.

