## Supplemental files for "Creatine phosphagen system regulates energy metabolism and growth of trophoblasts"

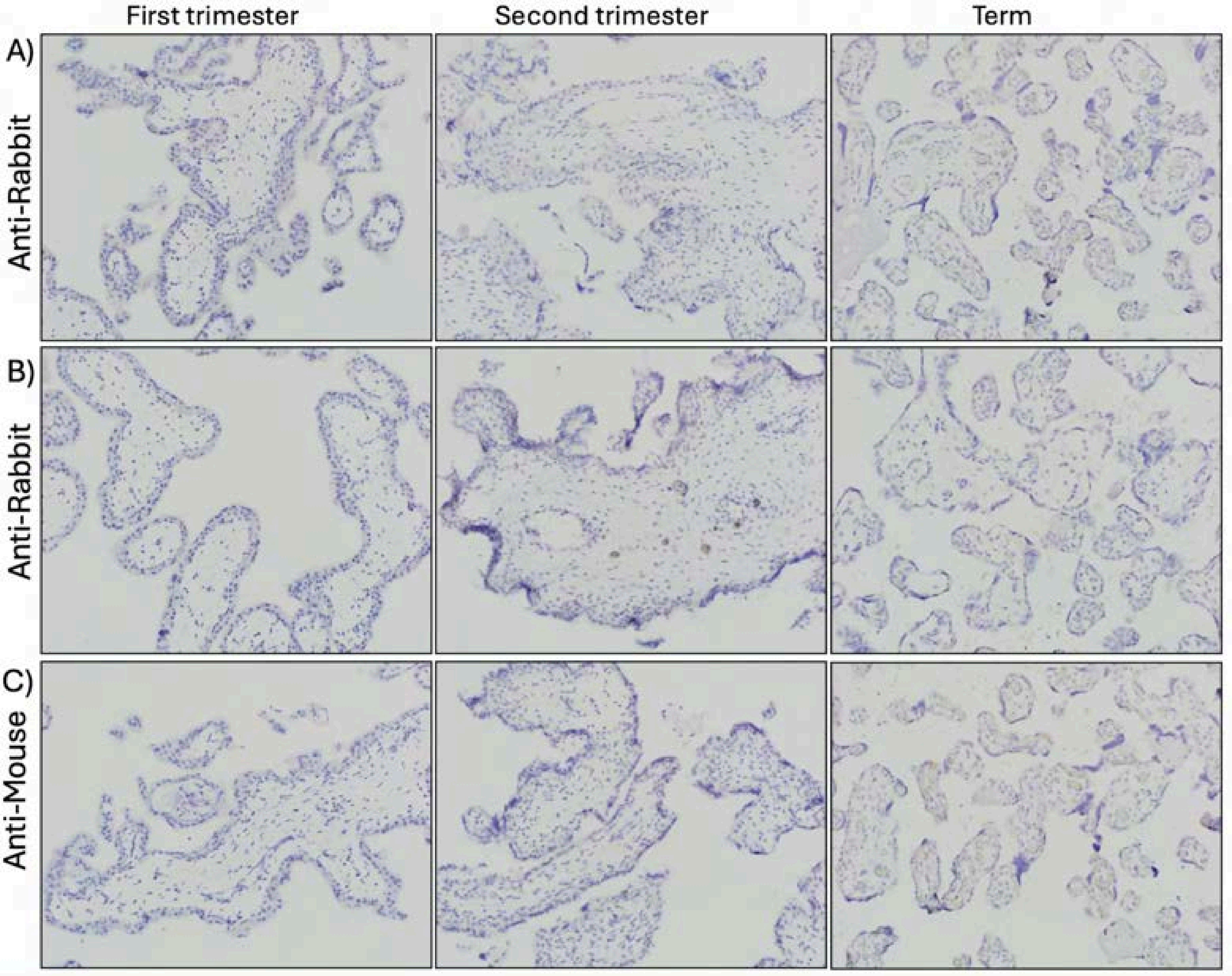


**Supplemental Figure S1**

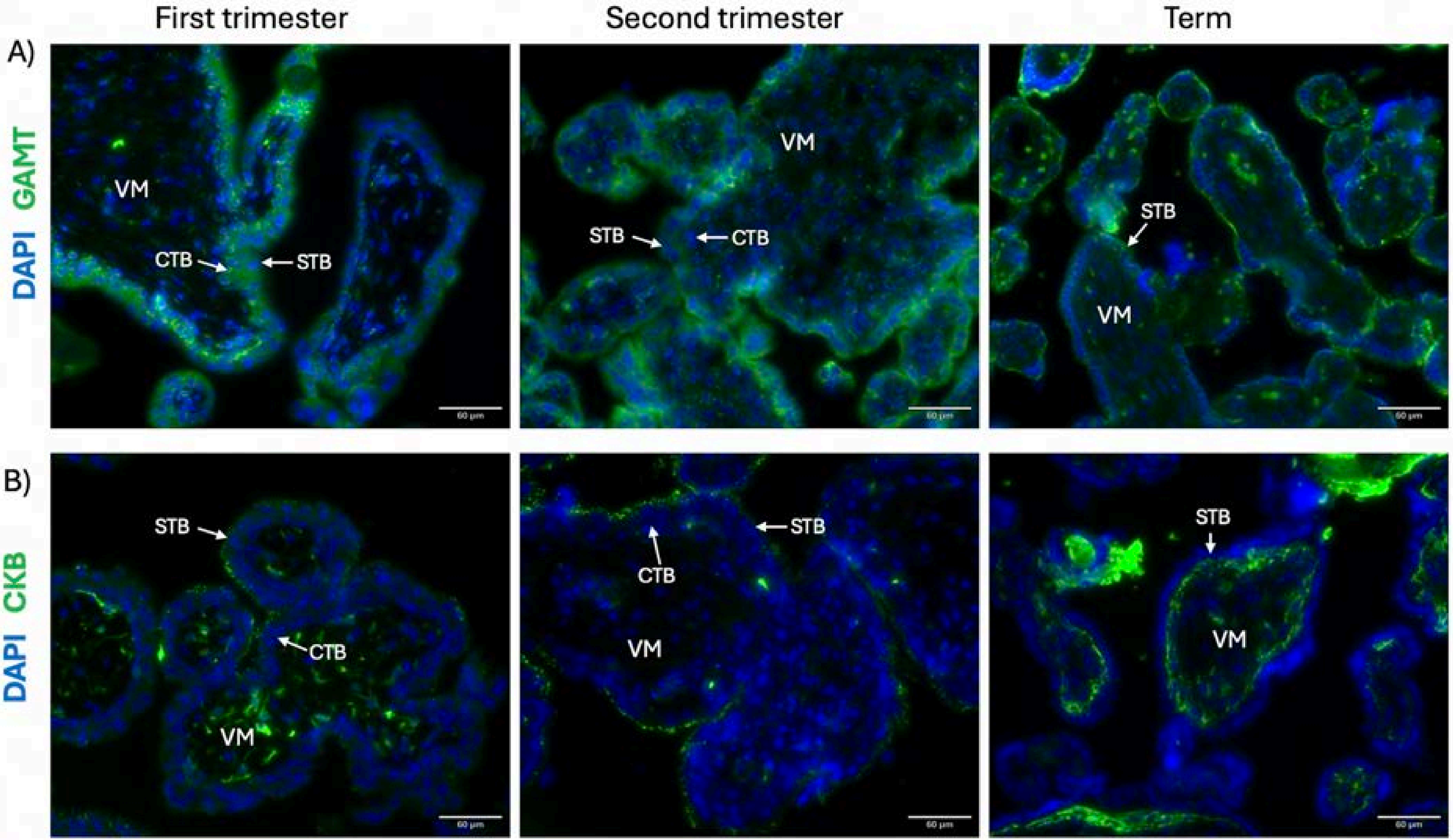


**Supplemental Figure S2**

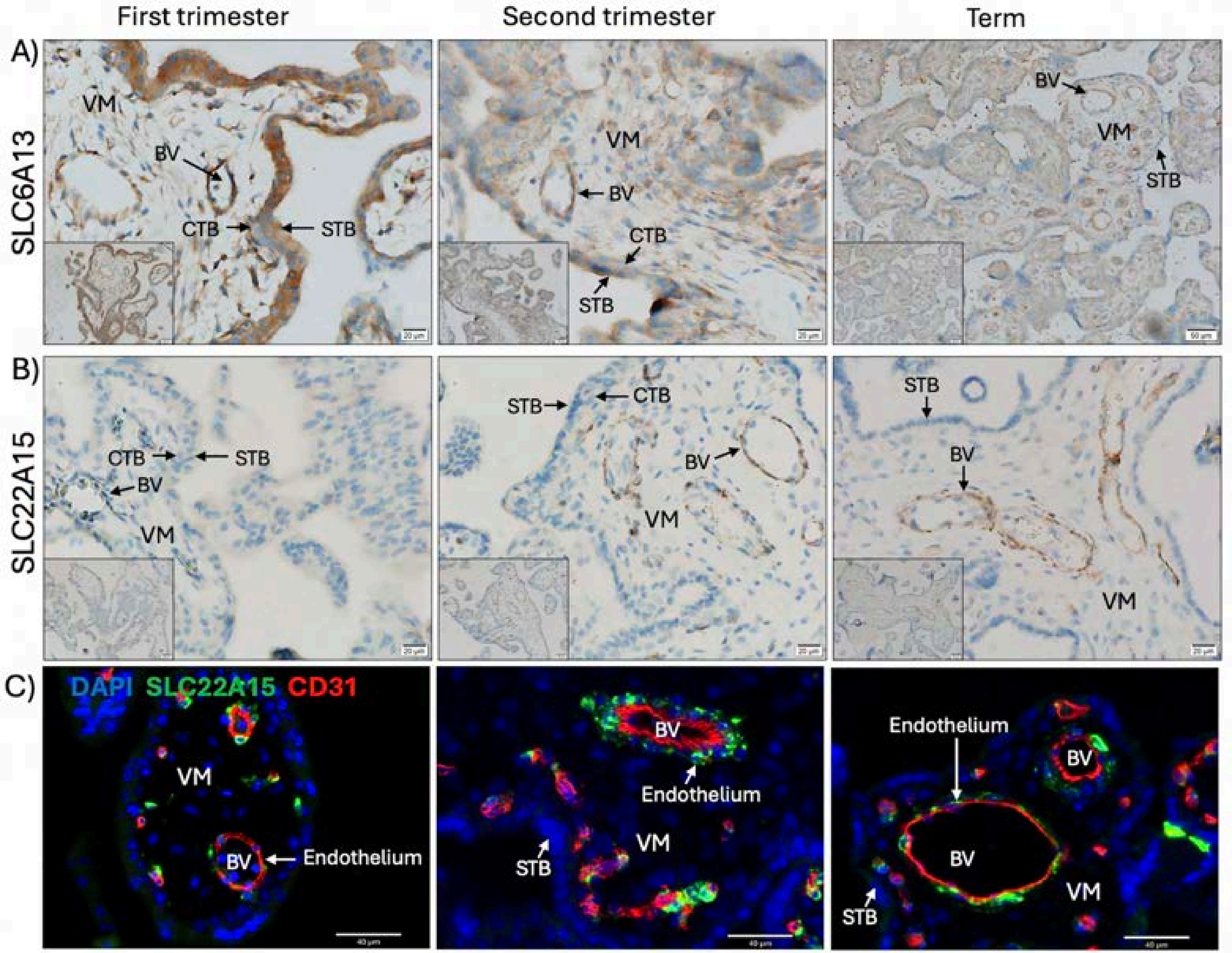


**Supplemental Figure S3**

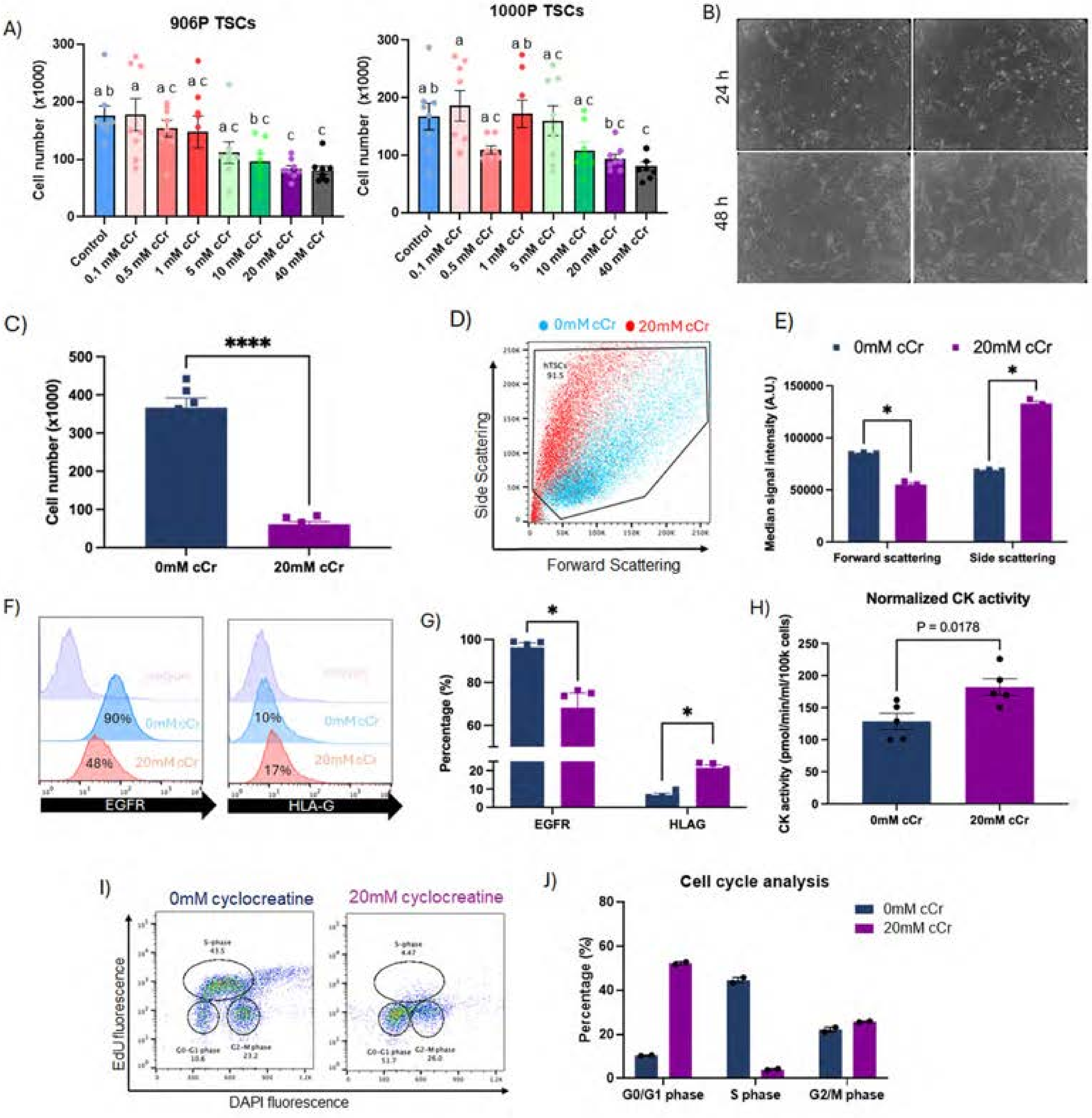


**Supplemental Figure S4**

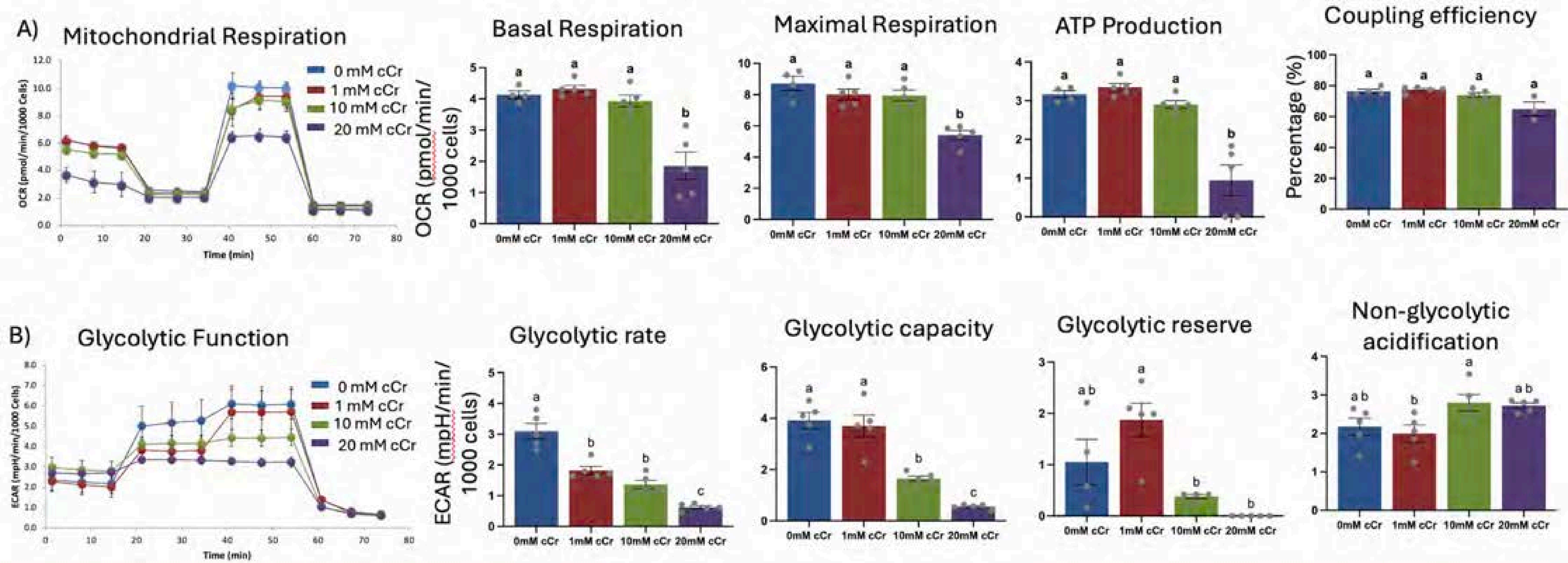


**Supplemental Figure S5**

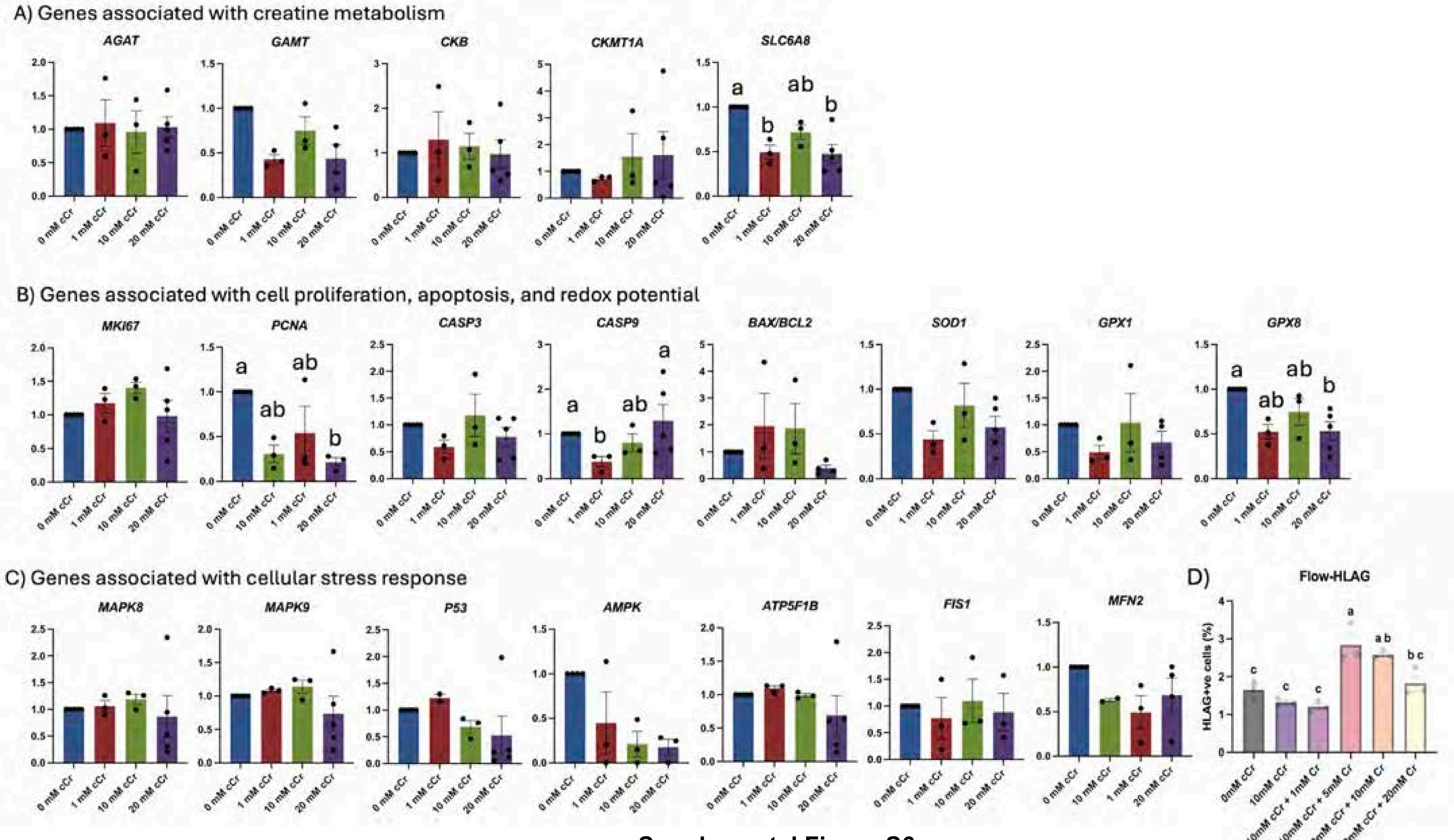


**Supplemental Figure S6**
